# Identification and Targeting of Regulators of SARS-CoV-2-Host Interactions in the Airway Epithelium

**DOI:** 10.1101/2024.10.11.617898

**Authors:** Brooke Dirvin, Heeju Noh, Lorenzo Tomassoni, Danting Cao, Yizhuo Zhou, Xiangyi Ke, Jun Qian, Sonia Jangra, Michael Schotsaert, Adolfo García-Sastre, Charles Karan, Andrea Califano, Wellington V. Cardoso

## Abstract

Although the impact of SARS-CoV-2 in the lung has been extensively studied, the molecular regulators and targets of the host-cell programs hijacked by the virus in distinct human airway epithelial cell populations remain poorly understood. This is in part ascribed to the use of nonprimary cell systems, overreliance on single-cell gene expression profiling that does not ultimately reflect protein activity, and bias toward the downstream effects rather than their mechanistic determinants. Here we address these issues by network-based analysis of single cell transcriptomic profiles of pathophysiologically relevant human adult basal, ciliated and secretory cells to identify master regulator (MR) protein modules controlling their SARS-CoV-2-mediated reprogramming. This uncovered chromatin remodeling, endosomal sorting, ubiquitin pathways, as well as proviral factors identified by CRISPR analyses as components of the host response collectively or selectively activated in these cells. Large-scale perturbation assays, using a clinically relevant drug library, identified 11 drugs able to invert the entire MR signature activated by SARS-CoV-2 in these cell types. Leveraging MR analysis and perturbational profiles of human primary cells represents a novel mechanism-based approach and resource that can be directly generalized to interrogate signatures of other airway conditions for drug prioritization.

## INTRODUCTION

The conducting airway epithelium is the first line of defense in the respiratory tract against pathogens and a prime site of SARS-CoV-2 viral entry, also serving to maintain a high viral load during lung infection. Single cell transcriptomics has been widely used to investigate viral-host interactions and to identify genes that may mediate interaction with viral proteins to support COVID-19 pathogenesis. Two main limitations have potentially affected the relevance of these studies. First, studies have initially explored non-physiologic cell lines sensitized to SARS-CoV-2 infection by transgenic ACE2-TMPRSS2 expression^1^. More critically, most studies have leveraged cell lines derived from non-lung sources or even cancer cell lines^2–8^. Although they uncovered key mechanisms of viral-induced host responses associated with deleterious cellular effects, they had limited impact in revealing tissue or cell type-specific disease mechanisms, especially as they relate to the highly heterogeneous lung and airways-specific epithelium. To address these challenges, recent studies have used human primary cells grown in organotypic cultures or human induced pluripotent stem cell (hiPS)-derived lung organoids, thus enhancing the relevance of these screens^9^.

Studies to elucidate the mechanisms by which SARS-CoV-2 hijacks the cellular function in the lung epithelium have relied largely on the analysis of single cell transcriptomic profiles. Since gene expression represents a poor proxy of protein activity, these results are intrinsically biased towards the downstream effects of virus-mediated host-cell reprogramming rather than the key proteins—including transcription factors (TFs) and co-factors—that represent their upstream mechanistic determinants. We and others have shown that these limitations can be effectively addressed by relying on network-based algorithms—such as VIPER^10^ and its single cell extension metaVIPER^11^—that can accurately identify Master Regulator (MR) proteins responsible for mechanistically controlling pathophysiologic gene expression signatures, via their transcriptional targets^12,13^. This is especially relevant at the single cell level where as many as 80% - 90% of genes produce virtually no reads, an effect known as gene dropout that significantly jeopardizes the ability to elucidate biological mechanisms from single cell data. Specifically, by analyzing the differential expression of a protein transcriptional targets—as identified by the extensively validated ARACNe algorithm^14^—VIPER assesses the contribution of every TF and co-TF to implementing a specific gene expression signature (henceforth ***protein activity*).** This allows for measuring the activity of proteins even when mRNA reads are minimally detected, making the algorithm especially well-suited to single cell analyses. VIPER has been shown to outperform antibody-based expression analysis in single cells, because (a) protein abundance is a suboptimal proxy for protein activity and (b) antibody availability and specificity is still limited. Indeed, VIPER has been extensively applied to investigate pathogenetic mechanisms in cancer leading to successful clinical trials^13,15–19,20,21^, as well as in non-cancer related fields, including immunology^22^, diabetes^23^, regenerative medicine^22,24^, neurodegenerative disease^25–27^, and stem cell biology^22,28,29^. Although we previously leveraged these network-based approaches to investigate SARS-CoV-2 ^30^, there were critical limitations. First, the analysis relied on cancer cell lines—a poor proxy for the understanding of the pathophysiologic responses of human lung epithelial cells; second, only the average effects of the virus on the multiple subpopulations that comprise the human lung epithelium were investigated, rather than the subpopulation-specific responses. Third, the library of drugs used in these analyses was limited to oncology drugs with relatively high toxicity.

Here we addressed these fundamental limitations by performing VIPER-based analyses of human airway cells, grown in organotypic cultures to identify the host-cell hijacking programs induced by SARS-CoV-2 in basal, ciliated, and secretory cells at the single cell level and at different stages of infection. Integration of VIPER-based MR activity with recent CRISPR-KO screens^3,31,32^ revealed both pan-airway epithelial and airway cell-type specific MR modules controlling the regulatory programs hijacked by SARS-CoV-2. Analysis of cells directly infected with SARS-CoV- 2 compared to bystander cells revealed non-interferon (IFN) signatures enriched in epigenetic regulators (histone deacetylases/DNMT), endosomal, ubiquitin and other pathways with components of these signatures identified in all or in specific cell types.

To identify small-molecule compounds capable of inverting these MR activities, we performed a large-scale perturbation screen in airway epithelial organotypic cultures using a library of FDA-approved drugs not limited to oncology. This screen identified 11 drugs able to target SARS-CoV- 2-mediated MR signatures across the three airways subpopulations. The network-based approach we used here (ViroTreat)^30^ allowed targeting the entire signature of MR proteins identified as mediators of the SARS-CoV-2 hijacking of the host cell programs, rather than an individual regulator. This is highly advantageous in a heterogeneous primary cell system, such as the airway epithelium, as the individual proteins controlling these effects may be different in different subpopulations.

Overall, the framework described here can be broadly extended to investigate mechanisms of viral-host interaction and MR-mediated reprogramming by a variety of other pathogens affecting the human airway epithelium as well as to prioritize clinically relevant drugs or novel agents as potential inhibitors. In particular, the drug perturbation assays—which represent the most expensive and time-consuming element of this study—provide a universal resource that can be leveraged to prioritize drugs targeting the host-cell MR signatures induced by virtually any other pathogen affecting lung epithelial cells, thus requiring only assessment of pathogen-specific gene expression signatures for MR analysis.

## RESULTS

### A Network-based approach identifies common and cell-type specific pathways of the SARS-CoV-2-induced host response in the human airway epithelium

To identify candidate regulators responsible of viral-induced hijacking of the host machinery and to investigate their corresponding biological functions, organotypic air-liquid interface (ALI) cultures from adult human extrapulmonary airways (lower trachea and main bronchi) were exposed to SARS-CoV-2 (MOI 0.1, *n* = 3, **Fig. 1a)** or mock conditions (see Methods) after being cultured for 21 days. Control and infected cultures were then analyzed at 1, 3, and 6 days post-infection (dpi). Infection efficiency was confirmed by plaque assays and by the identification of nucleocapsid (NP) signals in immunofluorescence (IF) assays (**Fig. 1a**).

**Figure 1:**
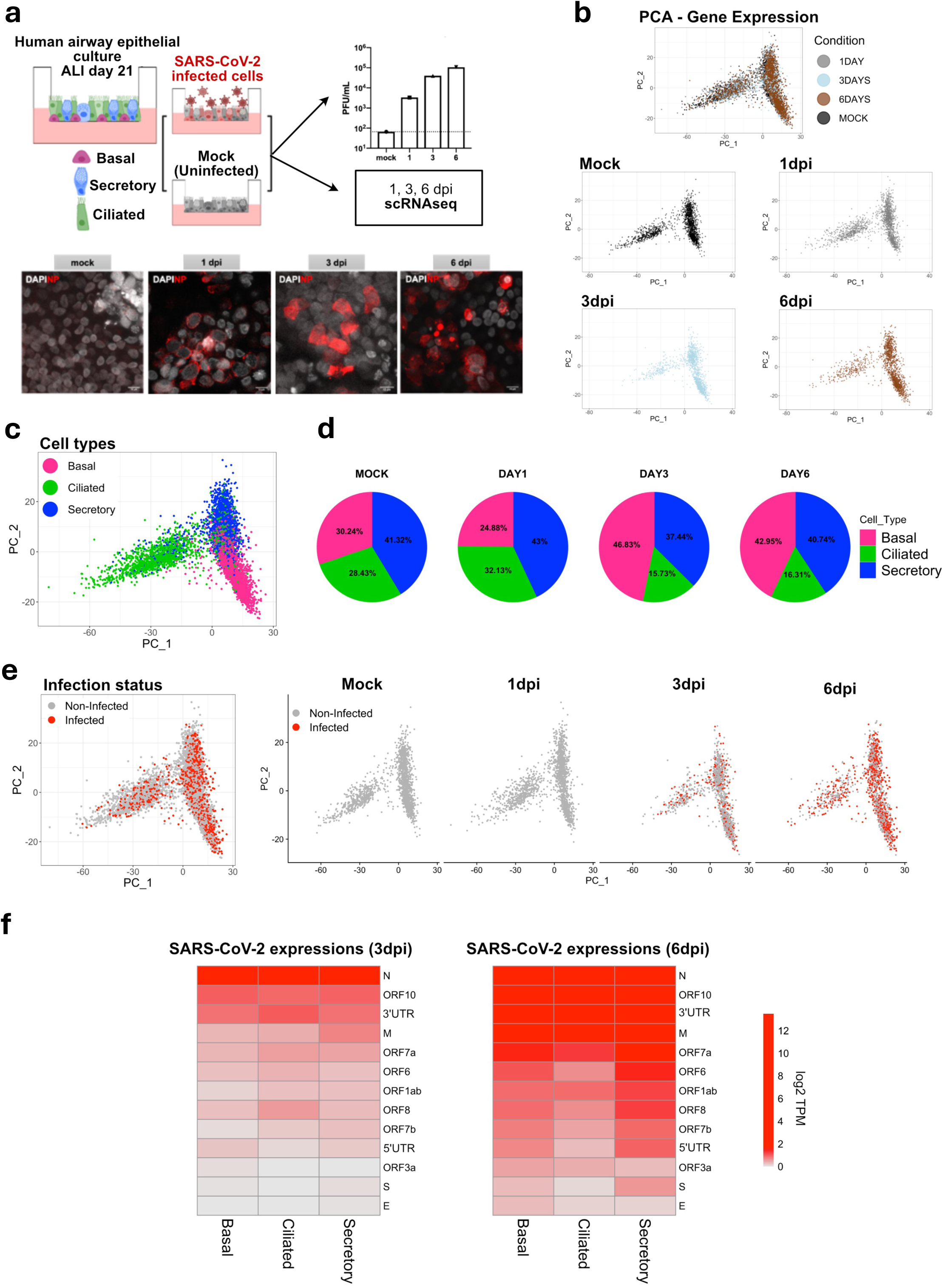
**SARS-CoV-2 infects distinct adult human airway epithelial cell populations in ALI organotypic culture** Diagram experimental design: human airway epithelial progenitor cells from adult donors cultured in Air-liquid interface (ALI) for 21 days were infected with SARS-CoV-2 (MOI: 0.1) or untreated (mock) and processed for scRNA-Seq at 1, 3, and 6 dpi. Graph: quantification of single stranded RNA (PFU/mL, n = 3 cultures per time point). Immunofluorescence (IF) and confocal imaging of cultures immunostained for the SARS-CoV-2 nucleocapsid (NP, red) protein and DAPI (gray). Scale bar = 11um. PCA plots based on gene expression data of quality control (QC) filtered cells. Each dot represents a single cell for all four different conditions (top panel). Cells are are also split into distinct sub-panels: MOCK (black), 1 dpi (gray), 3 dpi (light blue) and 6 dpi (brown). PCA plot based on Gene Expression data of QC- filtered cells. Cells are colored according to the different cell types: Basal (Pink), Ciliated (Green), and Secretory (Blue) Pie charts displaying the proportions of three cell types (basal, secretory and ciliated cells) in the scRNA-Seq data across time points (Mock, 1, 3, and 6 dpi). PCA plots based on Gene Expression data of infected (Red) and non-infected (Grey) cells from MOCK, 1 dpi, 3 dpi, and 6 dpi cultures. Infection status was determined by aligning raw data (FASTQ files) against the SARS-CoV-2 genome. Cells were considered infected if they contained at least one viral read mapped to the SARS-CoV-2 genome. SARS-CoV-2 components: Heatmap showing the average gene expression values (log- TPM) of reads aligned to the viral genome for each cell type at 3dpi and 6dpi.

Single-cell RNA-Seq (scRNA-Seq) profiles were generated and analyzed to identify proteins controlling the programs hijacked by the virus in basal, ciliated, and secretory cells, as well as to assess their effect on cell function. Quality control filtering (**Suppl. Table 1**) revealed minimal batch effect, with cells clustering according to the expected epithelial phenotypes, namely basal, secretory and ciliated cells, as assessed by analysis of established lineage markers and further validated via SingleR analysis^33^ using previously reported single-cell profiles.^34^ (**Fig.1b,c**; **Suppl. Fig. 1a-c**) .

Although the total number of epithelial cells was not significantly changed in the SARS-CoV-2- exposed and mock control cultures, the relative proportion of cell types differed over time in culture. The percentage of basal cells increased while that of ciliated cells decreased both at 3dpi and 6dpi, compared to controls (**Fig. 1d**, p ≤ 2.2×10^-16^, by Chi-Square test). Infection status was determined by alignment of quality-controlled cells against the SARS-Cov-2 genome confirmed by at least one read per cell mapped to the viral genome. Consistent with the single-stranded (ss) RNA results, the viral genome alignment analysis showed a progressive increase in the proportion of infected cells vs. non-infected cells at 3 and 6 days post-exposure (**Fig. 1e**). This also revealed *N* and *Orf10* among the most expressed viral genes in infected cells, across all cell types (**Fig. 1f**). In contrast to prior reports^34–36^, no statistically significant differences in infection rates were detected across these three cell populations (**Suppl. Table 2**), a trend that was independent of the specific threshold of the mapped reads to the SARS-CoV-2 genome used for the identification of infected cells (**Suppl. Fig. 1d**).

Next, we investigated the cell type-specific host response of ciliated, basal and secretory cells by analyzing the differential activity of all regulatory proteins, as assessed by metaVIPER^11^, the extension of VIPER to single-cell analyses. To generate sufficiently informative host response signatures for metaVIPER analysis, read counts were uniformly normalized using a metaCell approach (**Fig. 2a** see methods). MetaVIPER (*henceforth VIPER for simplicity*) was then used to transform the differential gene expression signatures of individual SARS-CoV-2-infected metaCells at 3 and 6 dpi—vs. metaCells from either mock controls or non-infected cells at the same time point—into differential protein activity profiles. For this purpose, we generated a context-specific regulatory network by analyzing a publicly available repository of primary human airway epithelial gene expression profiles^37,38^ with the ARACNe algorithm^10,11^. The results of this analysis was a subpopulation-specific repertoire of proteins that were aberrantly activated or inactivated in response to SARS-CoV-2 infection in basal, ciliated or secretory cells (**Fig. 2b, Suppl. Fig. 2a**).

**Figure 2:**
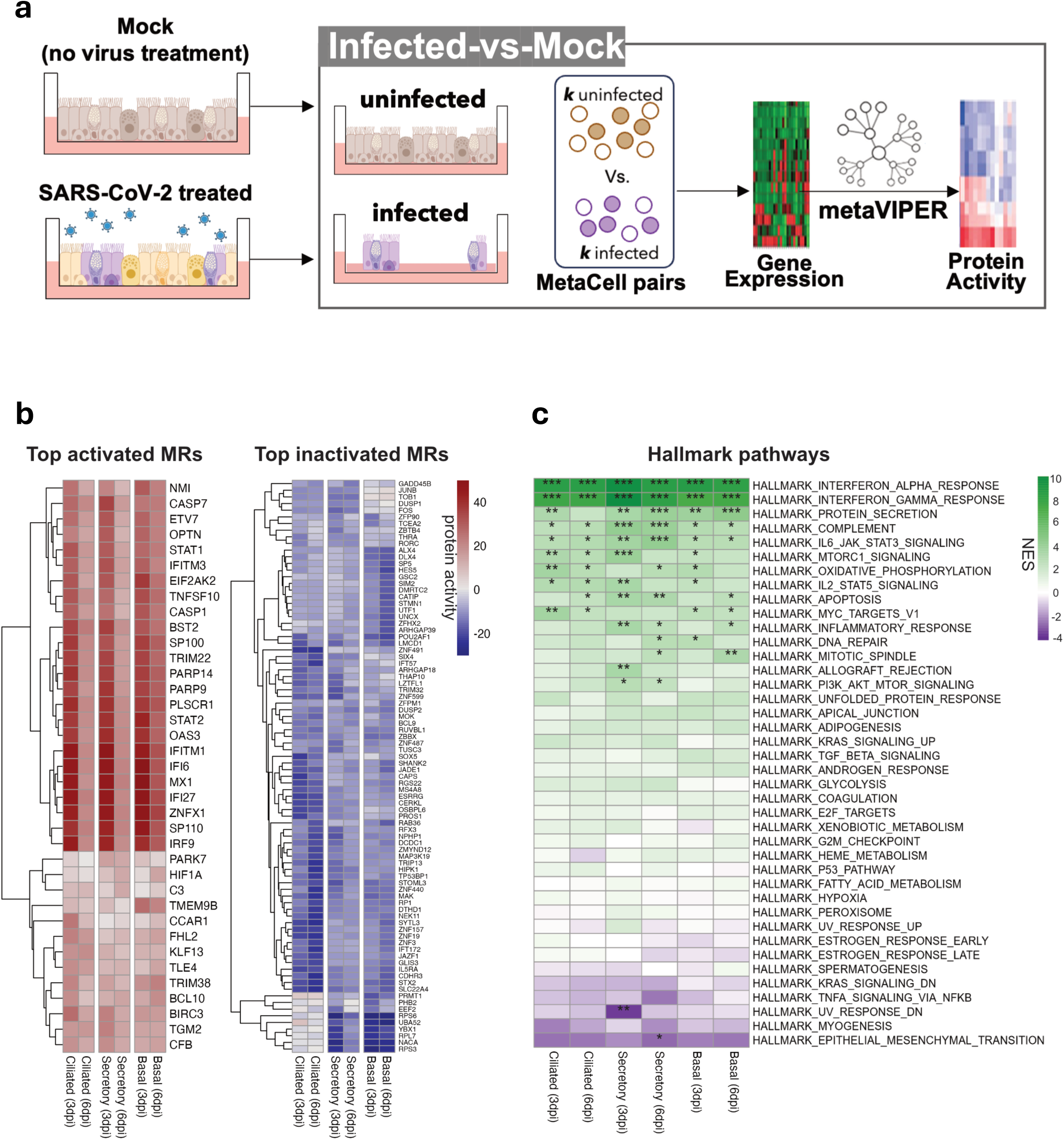
**Protein activity analysis identifies regulators and targets of SARS-CoV-2 infection in human airway epithelial cells.** Diagram: computational strategy for analysis of protein-activity and MRs of SARS-CoV-2- infection from *mock vs infected cultures (UIS: Unspecific Infection Signature*). Differential gene expression signatures of SARS-CoV-2-exposed (3dpi, 6dpi) vs mock (control) were generated for each cell type. Infected cells (n = *k*) vs mock cells (n = *k*) were randomly selected 100 times (i.e. repeated subsampling), then their gene expression signature was converted computationally into protein activity using VIPER (see Methods). The protein activity signatures obtained throughout 100 subsamplings were then integrated using Stouffer’s method for each cell type. Heatmap of the VIPER-inferred top differentially activated (left) and inactivated (right) proteins of SARS-CoV-2 *infected vs. mock* cultures in each time point (3 dpi, 6 dpi) and cell type (basal, secretory, ciliated). Top activated and inactivated proteins were retrieved from each cell type signature, and clustered, separately using hierarchical clustering algorithm (average-linkage method). Differential protein activity is shown as NES. Positive (brown) and negative (blue) values indicate protein activation and inactivation, respectively. Heatmap depicting differential enrichment of biological hallmark pathways in SARS-CoV- 2 *infected vs. mock* protein signatures at 3 and 6 dpi. NES was estimated by aREA (see methods). Enrichment of activated (purple) and inactivated (green) proteins is shown. Statistical significance indicated by asterisks: ***: p-value<0.001, **:p-value<0.01, *:p- value<0.05.

The rationale for comparing infected cells to either mock or non-infected cells is critical to deconvoluting unspecific/indirect effects (cytokine secretion by infected cells affecting the non-infected bystanders) from the specific/direct effects (the reprogramming unique to the infected cells, where the unspecific effects are subtracted). As such, we define the signature derived from infected vs. mock cells as the *Unspecific Infection Signature* (**UIS**) and the signature derived from infected vs. non-infected bystander cells as the *Specific Infection Signature* (**SIS**).

As expected, hallmark^39^ pathway analysis of the UIS signature revealed interferon alpha/gamma signaling, protein secretion, complement, and IL6/JAK/STAT3 signaling as the most significantly activated pathways in all cell types (**Fig. 2c, Fig. S2b**). Indeed, the 50 most differentially activated proteins included interferon-induced factors such as IFITM1—a known SARS-CoV-2 infection cofactor in the human lung^40^—IFI6, IFI27, MX1, IRF9, STAT1, STAT2, as well as a number of others proteins (ZNFX1, PLSCR1, SP110, PARP14, TNFSF10, CASP1, OAS3) that were prominently activated (*p* ≤ 0.001) in the host response signature (**Fig. 2b-c, Suppl. Fig. 2b, Suppl. Table 3a**)^41^.

When normalized enrichment scores (NES) were averaged over the three cell types at days 3 and 6, pathway enrichment analysis of the UIS signature—either based on differentially expressed genes or differentially active proteins—was highly concordant. Statistical significance of the concordance was assessed by Spearman correlation (ρ = 0.641, *p* = 8.2×10^-6^) and F-statistic (*F* = 62.0, *p* = 1.7×10^-9^). Results were also consistent with previous reports of significant innate immune response activation by SARS-CoV-2 in contexts as diverse as human airway-derived lung cancer cells (calu-3), gastrointestinal organoids of different origins^30^, and cultured human bronchial epithelial cell (HBEC) ALI cultures^34,39,40^ (**Suppl. Fig. 2c**).

Moving from pathway analysis to analysis of individual proteins, besides IFNs and related factors, the UIS also identified the zinc-finger protein ZNFX1, as a top activated MR in the three cell types analyzed, particularly at 3dpi (**Fig. 2b left**). Interestingly, activation of ZNFX1 in immune cells isolated from bronchoalveolar lavage of COVID-19 patients has been identified as an important component of the antiviral response^41^. Mechanistically, these studies show that ZNF proteins could directly recognize and bind to CpG sites in SARS-CoV-2 to induce IFN in infected cells. Thus, our findings of similar ZNFX1 activation in a system that consists solely of epithelial cells is noteworthy, suggesting that the ZNF-IFN association may occur in a broader cellular context. However, our MR analysis also revealed other ZNF family members among the top inactivated proteins in specific cell types. For example, ZNF491, ZNF157, ZNF19 emerged from the UIS as significantly inactivated in ciliated cells compared to secretory or basal cells (p ≤ 0.01, by t-test) (**Fig. 2b right**). Whether the differential inactivation of ZNF family members selectively in specific cell types influences the susceptibility to viral infection, remains to be investigated.

There is also increasing evidence that viruses may hijack specific ribosomal proteins to achieve optimal viral translation^42^. While broadly confirming these findings, our analysis also suggest that SARS-CoV-2 may selectively target the translational machinery in distinct cell types. For instance, we found specific ribosomal subunits (RPS6, RPS3, and RPL7)^43^, and translation-related proteins (NACA^44^, YBX1^45^, and PRMT1^46^) to be significantly inactivated only in secretory and basal cells (p < 0.01 by t-test) (**Fig. 2b right**).

### Analysis of infected vs bystander signatures reveal cell type-specific SARS-CoV-2 host response MRs

Previous studies show that IFN signaling from SARS-CoV-2 infected cells indirectly affects their immediate neighbor cells, thus contributing to a broad IFN signature activation across the entire airway epithelium^34^. To account for potential indirect, paracrine effects, such as those induced by IFN signaling, we reanalyzed our data, now comparing infected (≥1 viral RNA reads) with non-infected bystander cells (with no viral RNA reads) in SARS-CoV-2-treated cultures, as captured by the *Specific Infection Signature (SIS*). Specifically, we reasoned that analysis of the SIS signature could identify additional, more specific host cell response mechanisms that otherwise could not be revealed by the UIS, in which infected and mock cells were analyzed without bystander cell contribution.

For this, we focused on 3dpi to prevent inclusion of secondary effects (**Fig. 3a**). As expected, SIS analysis no longer highlighted “immune response” as differentially activated (**Fig. 3b-c** to **2b-c**). Instead SIS identified MRs activated specifically in the infected cells, in some cases also revealing cell-type specificity. For instance, hallmark features of SIS, such as protein secretion, apical junction, androgen response, and MYC target pathways were significantly activated in secretory cells, while mitotic spindle was found activated mostly in ciliated and secretory cells. Hallmark estrogen response was found in basal and ciliated cells but not in secretory cells (**Fig. 3c, Suppl. Fig. 2b, Suppl. Table 3b**). Viral infection is facilitated and propagated by heightened cell to cell transmission in apical junctions of the luminal mucociliary cells, with virion accumulation in the mucus layer, facilitating infection of neighboring cells^47–49^. SARS-CoV-2 has also been proposed to hijack microtubule kinases crucial for organization of mitotic spindle to influence viral load and invasion. Based on this, anti-cancer drugs that block mitosis through disruption of microtubule organizing center have been tested in COVID-19 human clinical trials^50^. Yet, since these effects appear to be largely population cell-specific, these drugs may not be effective in an heterogeneous cellular context, thus underlining the relevance of the information provided by the SIS. The seemingly unrelated pathways described above collectively reflect the complex orchestration of events associated with COVID pathogenesis in the airway epithelium, a system known for its significant cellular diversity.

**Figure 3:**
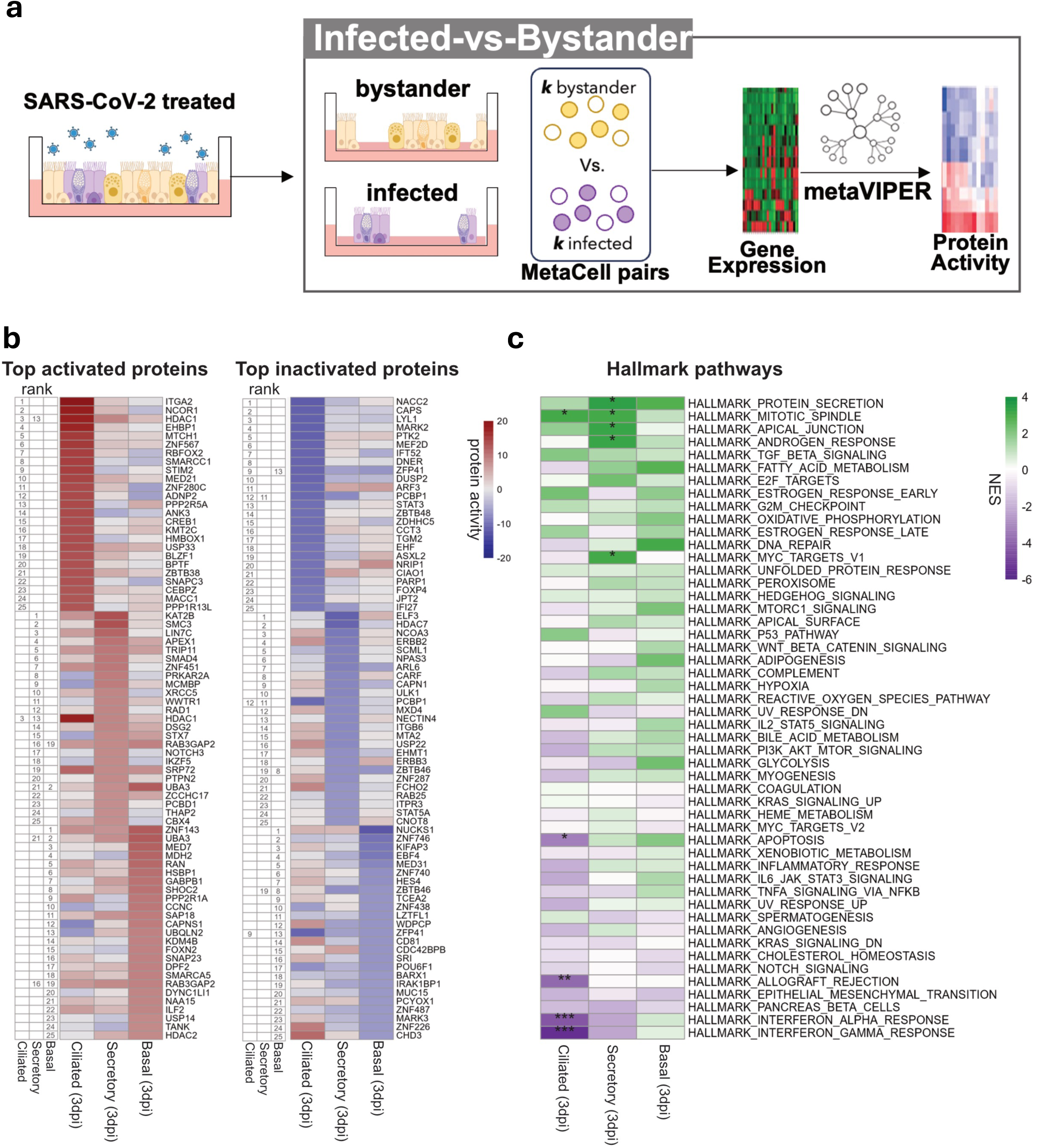
Protein activity analysis identifies the top master regulators of the host response in distinct populations of human airway epithelial cells infected with SARS-CoV-2 compared to bystander cells (SIS*: Specific Infection Signature*). Diagram computational strategy: analysis of protein-activity and MRs *in infected vs. bystander* cells from SARS-CoV-2-treated cultures at 3 dpi. Infected (n = k) vs bystander (n = *k*) cells were randomly selected 100 times (as before), then gene expression was converted into protein activity (VIPER, see Methods). The protein activity signatures obtained throughout 100 subsamplings were then integrated using Stouffer’s method for each cell type. Heatmap of the VIPER-inferred top 25 differentially activated (left) and inactivated (right) proteins of SARS-CoV-2 infected vs. bystander cultures at 3 dpi for each cell type analyzed. Ranking is shown numerically on the left of each heatmap. Differential protein activity is shown as NES. Positive (brown) and negative (blue) values indicate protein activation and inactivation, respectively. Differential enrichment of biological hallmark pathways in SARS-CoV-2 *infected vs. bystander* protein signatures at 3 dpi. NES was estimated by aREA (see methods); activated (purple) and inactivated (green) proteins are shown. Statistical significance: ***: p-value<0.001, **:p-value<0.01, *:p-value<0.05

At the individual protein level, analysis of the SIS signature identified ITGA2, NCOR1 and HDAC1 as the top three most activated host-cell proteins in ciliated cells following SARS-CoV-2 infection (**Fig. 3b**). ITGA2 encodes the alpha subunit of a transmembrane receptor integrin binding present in anchoring junctions to mediate cell-cell and cell-matrix interactions. A number of studies show that human adenovirus, herpesvirus, and other viral pathogens internalize into cells through a mechanism of integrin-mediated endocytosis^51,52^. Notably, there is evidence of SARS-CoV-2 binding to Integrin and activation of endocytosis as an essential ACE2-independent component of the viral infection^51,53^. ITGA2 has also been linked to p38 and p16-mediated senescence of SARS-CoV-2-infected cells^54^.

The two additional MR proteins, representing key putative mechanistic determinants of SARS-CoV-2 infection in ciliated cells, were NCOR1 and HDAC1. NCOR1 is part of a family of transcriptional co-repressors shown to form complexes with both class I and class II histone deacetylases (HDAC) in vitro and in vivo to repress gene expression. HDAC inhibitors repress ACE2 expression and other genes responsible for viral entry. Yet, the specific mechanisms by which activation of NCOR-HDAC in ciliated cells promotes viral entry are not fully understood^55,56^. Other HDAC family members identified in our screen included HDAC7 and HDAC2, among the most activated proteins in secretory cells and basal cells, respectively (**Fig. 3b**). Of interest, HDAC2 has been shown to interact with NSP5, the main SARS-CoV-2 protease to inhibit HDAC2-mediated IFN responses^2^. Our finding of the lysine acetyl transferase KAT2B as the most aberrantly activated MR in infected secretory cells could further support the idea of a mechanism of SARS-CoV-2 repression of IFN-related gene expression via chromatin remodeling to increased virulence. SMC3 (Structural Maintenance of Chromosomes 3) emerged as the second most aberrantly activated MR in secretory cells. SMC3 is a key component of the cohesin complex, which plays a critical role in the regulation of chromosome structure and gene expression. SMC3 is reported as a non-histone substrate of HDAC in cancer and has been shown to be involved in the restructuring of the chromatin architecture in SARS-CoV-2 infection^57,58^. Specific activation of these mechanisms in infected secretory cells is intriguing and worth further future studies.

Consistent with prior studies, VIPER analysis identified MRs representing key, complementary subunits of the middle module of the mediator complex, including MED21 and MED7—aberrantly activated in ciliated and basal cells, respectively—and MED31—aberrantly inactivated in basal cells. This complex associates directly with RNA polymerase II to regulate its function^59^. Biochemical structural data have shown that MED7 and MED21 form a hinge that enhances stable interactions between RNA polymerase II and the mediator complex, indicating that the activation of these proteins may be associated with increased transcriptional machinery activity^59^. Together the data suggested engagement of distinct SARS-CoV-2-mediated epigenetic reprogramming mechanisms in these populations of airway epithelial cells.

### Proviral host factors crucial for SARS-CoV-2 infection are selectively induced in different cell types

We then examined whether host factors previously reported to physically interact with SARS-CoV-2 proteins or shown to be critical for infection (proviral replication), based on CRISPR studies^3,31,32^, were differentially enriched in MR proteins. Host factors co-opted by SARS-CoV-2 during infection have been previously investigated, using genome-scale CRISPR knockout screens, in multiple cell types including Vero.E6 (monkey)^6^, Huh.7.5 (human hepatocarcinoma)^4^, A549 (human lung adenocarcinoma)^32^, and Calu-3 (immortalized human airway-like cell line derived from submucosal glands)^3,31^. These screens helped identifying several genes, including ACE2 and HMGB1, whose inactivation increased the viability of infected cells compared to non-targeting sgRNAs. From these studies, we selected the top 50 TFs and cofactors identified by CRISPR screens in A549 and Calu-3, based on their common origin from the lung/airway epithelium.

Indeed, UIS analysis recapitulated several of these proviral factors among the 50 most differentially active proteins (i.e., candidate MRs of host-cell hijacking, *hereafter MRs for simplicity*) across all cell types, at both 3 dpi and 6dpi (**Fig. 4a**). Leading-edge analysis revealed IRF1, IRF9, and STAT1 as the top most conserved factors across all three cell populations (**Fig. 4b**). The identification of interferon-regulatory factors and core transcriptional regulators of the inflammatory response as leading edge proteins—i.e., proteins identified among the most significant MRs but also positively modulating virulence, as supported by previously published CRISPR screens^3,31,32^—was intriguing as it suggested a dual role for these proteins as both proviral and antiviral factors during SARS-Cov-2 infection. Additional top leading-edge proviral factors identified by UIS signature analysis included MRs of distinct functional classes, including several previously implicated in viral internalization. Among these, four ATPase and accessory proteins family members (ATB8B, ATP6AP2, ATP6V1C1, ATP6V1H) were among the most aberrantly activated. Studies using bafilomycin to inhibit vacuolar-ATPase^60^ suggest that these ATPases act as proviral factors by facilitating SARS-CoV-2 entry through the endosomal route. This is further supported by our findings of the kinesin KIF13B and the small GTPAse RAB14, known regulators of intracellular trafficking, vesicle formation and endosomal recycling^61,62^, as key leading edge MRs (**Fig. 4b**).

**Figure 4.**
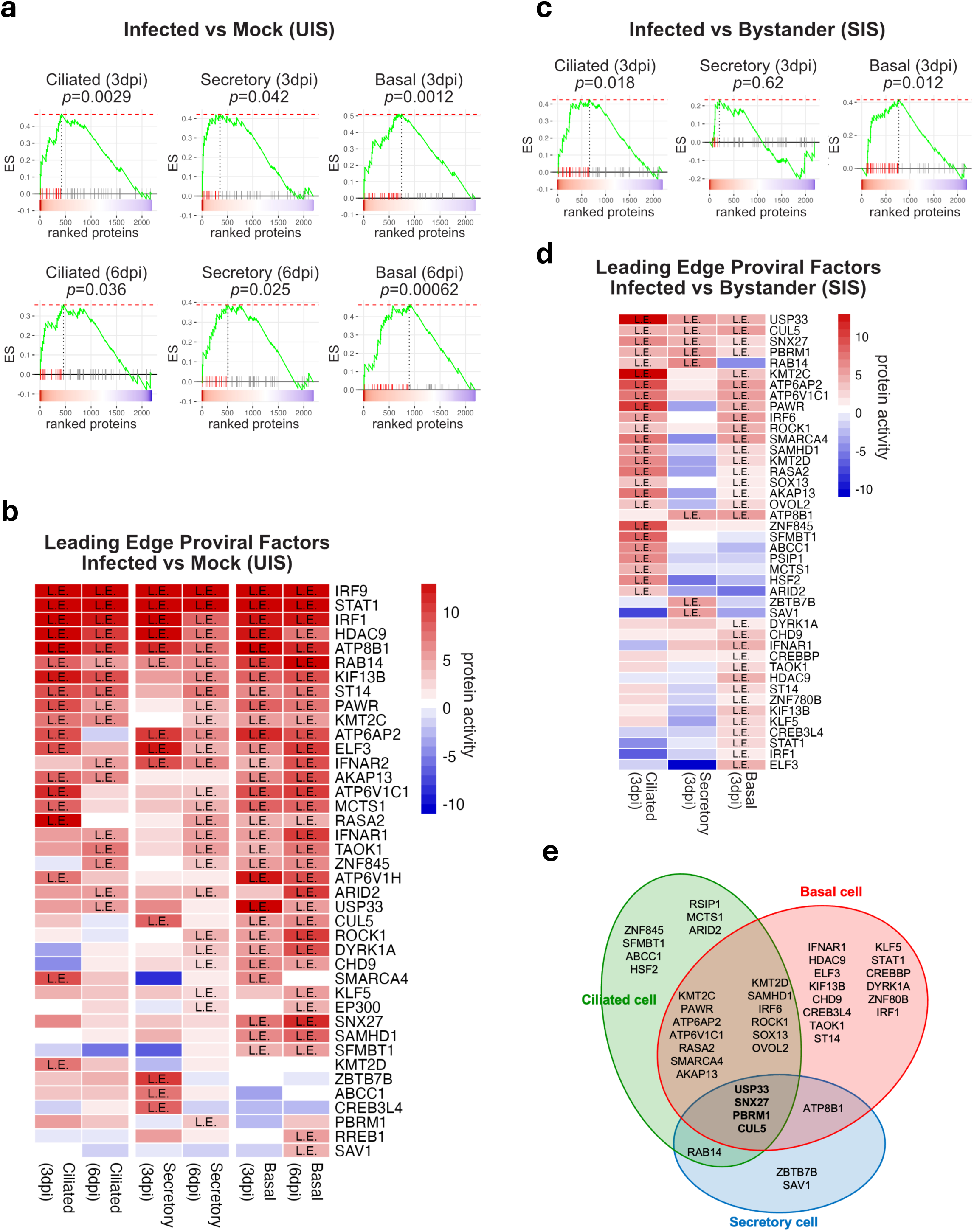
**Proviral host factors crucial for SARS-CoV-2 infection are induced in different airway epithelial cell types** Enrichment plots of proviral factors (n=50) in the SARS-CoV-2-*infected vs. mock* host response signature at 3dpi and 6dpi in each cell type. The color map on the bottom of each condition represents the host response signature (red; activated, blue; inactivated). Each bar represents the ranking enrichment of one proviral factor retrieved from previously identified gene sets from CRISPR-KO assays in human lung cancer cell lines (A549 and Calu-3). The green line represents the accumulative enrichment score. The red dotted line (top) shows the maximum enrichment score reached in GSEA. Hits on the left side of the vertical dotted line represent the subset of leading edge proviral factors among all 50 found in each condition. Enrichment p-values were calculated by fGSEA^84^. Heatmap of protein activity (red: positive; blue: negative) of proviral factors found in the leading edge (L.E.) of SARS-CoV-2 *infected vs. mock* GSEA plots from each condition. Rows were sorted in decreasing order of L.E. frequency across signatures Enrichment plots of proviral factors (n = 50) in the SARS-CoV-2-*infected vs. bystander* host response signature at 3dpi in each cell type. p-values were calculated by fGSEA and Leading-edge subsets were represented as in A. Heatmap of protein activity of proviral factors at the leading edge (L.E.) of SARS-CoV-2 *infected vs. bystander* GSEA plots in each cell type at 3dpi. Rows sorted in decreasing order of L.E. frequency across signatures. Venn diagrams summarizing the overlap of leading-edge host factors identified in A among the three cell types at 3dpi.

We then examined which of the top 50 proviral factors were also nominated as MRs by the SIS analysis. USP33, CUL5, SNX27, and PBRM1 emerged as the only leading-edge proviral factors identified as activated MRs in all the three cell types. (**Fig. 4c-e**). Interestingly, two of these factors have been functionally implicated in the regulation of ubiquitination in viral infection. USP33 is a deubiquitinase enzyme reported to modulate the host inflammatory response and antiviral activity through regulation of the turnover of IRF9^63^. CUL5, a component of the Cullin-5-RING E3 ubiquitin-protein ligase complex, has been shown to act as a critical antiviral host factor promoting ubiquitin-dependent degradation of the SARS-CoV-2 ORF9b protein^64^. The Sorting Nexin27 protein (SNX27) controls cargo recycling from endosomes to the cell surface inhibit viral lysosome/late endosome entry by regulating ACE2 abundance. Indeed, SNX27 can be hijacked by SARS-CoV-2 to facilitate viral entry^65^. PBRM1 is a chromatin modulator of the SWI/SNF chromatin remodeling complex, which includes *SMARCA4*, *SMARCB1*, *SMARCC1, ARID1A*, *DPF2*, *SMARCE1,* also implicated in *ACE2* expression regulation and, thus, in host cell susceptibility to viral infection^66^.

Taken together, these data reinforce the hypothesis that epigenetic reprogramming represents as a key mechanism leveraged by SARS-CoV-2 for hijacking host-cell programs in the airways epithelium. The analysis also helped identifying proviral factors that are either preferentially activated in a specific SARS-CoV-2-infected subpopulation or those shared across different subpopulations. Intriguingly, among the 50 leading-edge proviral factors present in the SIS signature, most were found activated in ciliated, basal cells or in both populations but not in secretory cells (**Fig. 4d-e**). Comparative analysis of the SIS and UIS MR signatures confirmed a lower number of proviral factors differentially activated in secretory cells (**Suppl. Fig. 3**). However, it is important to note that SARS-CoV-2 induces a robust MR activation selectively in secretory cells, as demonstrated by the SIS analysis of this subpopulation (**Fig. 3b**).

Taken together, the fact that our analysis identified key proteins known to modulate SARS-CoV- 2 infection, suggests that top population cell-specific MRs, as well as those conserved across multiple cell populations, that were not previously characterized may represent key regulators of novel mechanisms of host-cell hijacking by the virus, for future low-throughput validation assays. Moreover, the diversity of factors revealed by these analyses further reinforced the idea that the signature elicited in the airway epithelium by SARS-CoV-2 represents a complex combination of host defense and virus-hijacked signals.

### A large-scale functional drug screen identifies candidate drugs that effectively invert the SARS-CoV-2-induced MR signatures in the human airway epithelium

To identify drugs capable of inverting the MR activity signature induced by SARS-CoV-2 infection, we performed a large-scale drug perturbation screen in these organotypic airway epithelial cell cultures, thus supporting prediction of candidate MR-inverter drugs using the *ViroTarget* and *ViroTreat*^30^ algorithms. These represent the direct extension to the viral infection context of OncoTarget^18,20^ and OncoTreat^18,19,67^, two CLIA-compliant (https://www.cms.gov/medicare/quality/clinical-laboratory-improvement-amendments) algorithms extensively validated in both a pre-clinical and clinical oncology contexts. Drugs predicted as MR activity inverters by OncoTreat, from both bulk^18,67^ and single-cell^68^ profile data, have been validated by rigorous *in vitro* and *in vivo* assays. Consistent with this premise, we proposed that pharmacologic targeting of either individual MRs (ViroTarget) or of the entire MR protein module that regulates the virus-induced host-cell response (ViroTreat) could mitigate viral replication and infection co-morbidity. Briefly, ViroTarget analyzes the list of MRs to assess whether any of them may represent a high-affinity binding target of a clinically-relevant drug, among the 1,738 in DrugBank^69^. In contrast, ViroTreat assesses inversion of the virus-induced MR activity signature by assessing the enrichment of the 25 most activated and 25 most inactivated viral infection MRs in proteins differentially inhibited and activated in drug vs. vehicle control-treated cells, respectively. For this analysis, we used the 3dpi SIS signature, representing the most specific early hijacking of host cell programs by the virus.

ViroTarget identified 32 MRs (p ≤ 0.05, Benjamini-Hochberg corrected) aberrantly activated in at least one of the infected cell types (ciliated, basal, or secretory cells)—as high-affinity targets of 87 small molecule inhibitors in DrugBank. (**Suppl. Fig. 4**). Druggable proteins associated with epigenetic control of gene expression, such as the HDAC family members HDAC1, HDAC2, HDAC3 and HDAC9, were identified as SIS MRs in distinct cell types. Consistently, HDAC inhibitors, such as romidepsin identified by Virotarget were already proposed as antiviral drug candidates^30,70^. Additional MRs representing chromatin remodeling enzymes include EZH2, DNMT1 and CHD1; the latter is a chromatin organization modifier implicated in the recruitment of Influenza virus polymerase to promote viral multiplication^71^. These findings underscore the relevance and need for further investigation of epigenetic mechanisms hijacked by SARS-CoV-2 to induce host-cell reprogramming. ViroTarget also identified two out of three RXR retinoid receptors as candidate druggable MRs, as also supported by independent evidence from the functional analysis of SARS-CoV-2 infected human IPSC-derived organoids^9^. Another notable druggable MR identified by ViroTarget was KRAS. This protooncogene found frequently mutated in human cancers has been implicated in viral stress responses mediated by GRP78 a chaperone induced by SARS-CoV-2 infection^72,73^. Taken together, these data suggest that ViroTarget analysis can recapitulate several previously identified drugs as well as nominate additional ones for follow-up validation.

ViroTreat analysis requires a large-scale compendium of RNA-Seq profiles representing the response of cells to treatment with a large library of clinically relevant compounds. For this purpose, we used the PLATE-Seq technology and VIPER, developed by our labs^10,11,74^ to generate protein activity from the RNA-Seq profiles of airway epithelial cultures treated with 441 FDA-approved drugs. We selected drugs with well-characterized bioactivity, safety, and bioavailability properties, as determined by preclinical and clinical studies. VIPER analysis of RNA-Seq profiles of drug vs. vehicle control-treated cells helps characterize the proteome-wide mechanism of action (MoA) of each drug, which can be used to assess the drug’s ability to invert the activity of the MR signatures identified from VIPER analysis of the SIS signature of each individual subpopulation (**Fig. 5a**) (see methods). Here we define MoA as the repertoire of proteins that are differentially activated or inhibited by a drug in a tissue of interest, including high-affinity binding targets, secondary lower-affinity targets, and context-specific indirect targets. Taken together, these proteins define the drug’s pharmacologic activity^18^.

**Fig 5:**
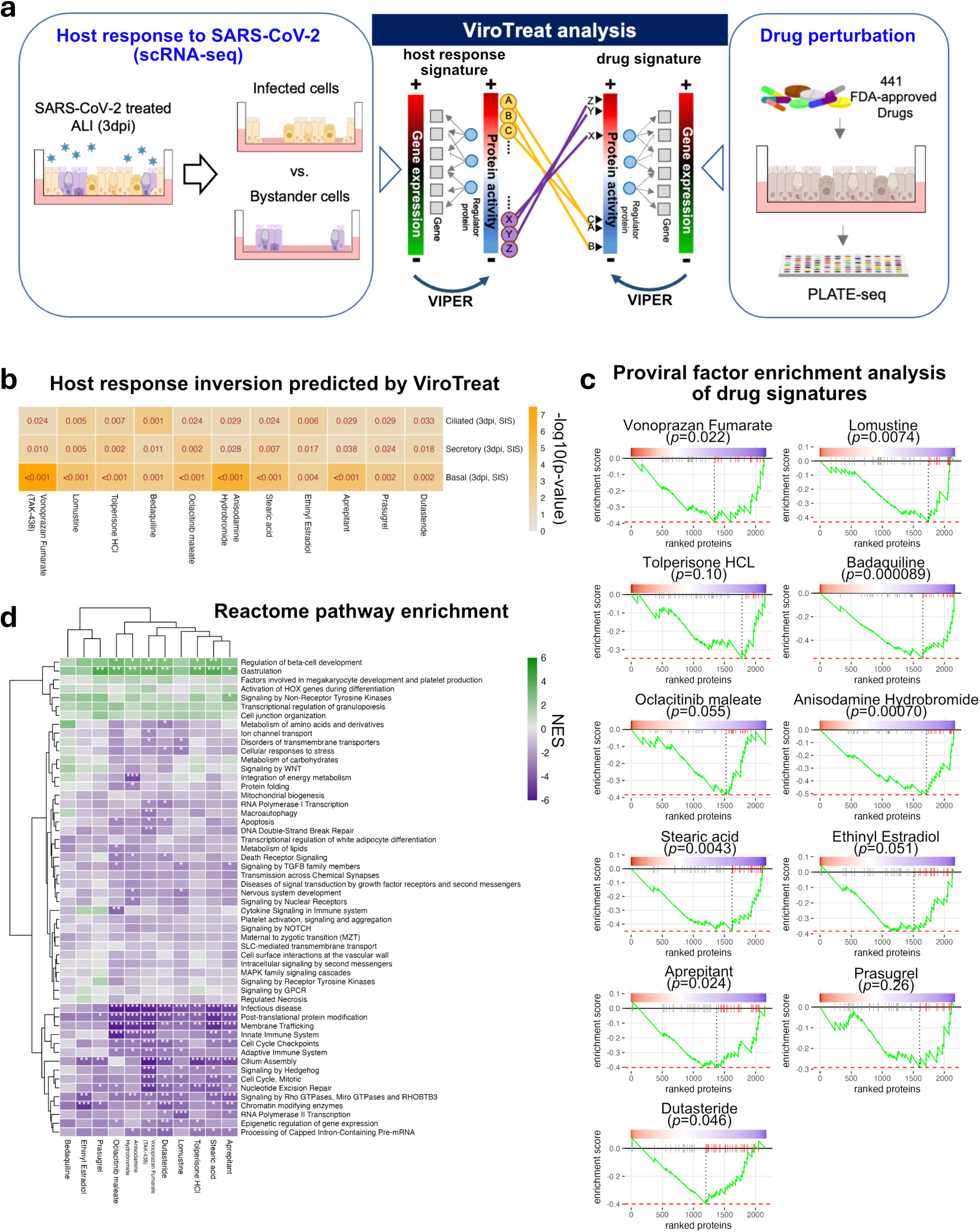
P**h**armacologically **targeting regulators of SARS-CoV-2-host interactions in the Airway Epithelium** Schematic representation of the *ViroTreat* workflow approach. A high-throughput drug perturbation screen was performed by treating ALI day 21 human airway epithelial progenitor cells with 441 FDA-approved drugs. Gene expression signatures were determined by PLATE-seq, converted to protein activity using VIPER and compared to the signatures previously obtained from SARS-CoV-2 *infected vs. bystander* cells. The Virotreat algorithm was employed in these paralleled signatures to identify candidate drugs predicted to invert the activated (yellow) or deactivated (purple) host response signature of infected cells. Heatmap of statistical significance (-log10 p-value) for the top 11 drugs predicted by ViroTreat as strong inverters of the SARS-CoV-2 host response signatures from infected vs. bystander cells at 3 dpi. The smaller p-value corresponds to a stronger inverter of the host response. Gene set enrichment plots for drug signatures. The gene set consisted of proviral factors which were included in the leading-edge subsets for host responses at least for one cell type (n = 42). The negative enrichment score curve (green) indicates that the proviral set was negatively enriched in the drug signatures able to invert the host response signatures Heatmap representing the reactome pathway enrichments (NES) in the signature of the top 11 drugs. The positive (green) and negative (purple) NES indicates the activated and inactivated pathways. Rows and Columns of the heatmap were sorted based on a complete-linkage hierarchical clustering method. Asterisks depict: *p<0.05, **p<0.01, ***p<0.001

For this purpose, drugs were added to differentiated ALI day 21 airway organotypic cultures at the their C_max_ concentration and RNA-Seq profiles were generated at 24h post treatment, using the fully automated PLATE-Seq technology^74^ (see methods) and analyzed by VIPER. Out of 441 drugs, ViroTreat identified 11 as statistically significant MR-inverters across all three airway epithelial cell types (**Fig. 5b, Suppl. Fig. 5a,b, Suppl. Table 4**). We further investigated the enrichment of MR in the leading edge of prior CRISPR screens, as described in the previous section (**Fig. 4d,e**). ViroTreat analysis showed leading edge MR activity inversion in nearly all cell types, with 9 out of 11 drugs (**Fig. 5c, Suppl. Fig. 6a**). Notably, bedaquiline—an established inhibitor of ATP synthase in drug-resistant pulmonary tuberculosis^75^—was identified as a candidate inverter of USP33 and CUL5 activity.

Pathway enrichment analysis using gene sets from Reactome Pathway (RP)^76^ showed that 8 out of the 11 drugs identified by ViroTreat also induced statistically significant negative NES for pathways associated with virus-induced processes, including membrane trafficking, infectious diseases, post-translational protein modification, and Rho GTPase signaling (**Fig. 5d**). Among the MRs whose activity was inverted by these drugs we found SMC3 and MED7 (see previous sections), as well as CREB1, USP33, ZFK451, SMARCA5 and others proteins reported in SARS-CoV-2-host interactions and pathogenesis databases (**Suppl. Fig. 6b**)^61^. ViroTreat also identified USP33, PAWR, ATP6AP2, CUL5, ROCK1, RAB14 as proviral MRs inactivated by 11 out of the 411 drugs screened. While most of the factors targeted were enriched in ciliated cells, two of them (CUL5, USP33) were present in all cell types (**Suppl. Fig. 6a**). Thus, the effect of ViroTreat- inferred MR-inverters drugs extends to proteins controlling mechanisms associated with viral replication. As such, these drugs should not only mitigate the hijacking of host-cell programs but also directly restrict viral replication.

## DISCUSSION

Here we presented a multi-pronged approach to elucidating the MR proteins representing mechanistic determinants of SARS-CoV-2-mediated hijacking of host cell mechanisms, as well as to identifying the drugs that can invert their activity. This highly generalizable approach integrates single-cell network-based analyses and large-scale drug perturbation assays in physiologically relevant air-liquid interface organotypic cultures of human airway epithelial cells representing the distinct subpopulations that comprise the airways epithelium. The study recapitulates key mechanisms identified in previous studies and nominates critical novel, population cell-specific MR proteins controlling host-cell hijacking by the virus to improve replication and viral life cycle (ViroCheckPoint) as well as the drugs that can inhibit their activity for follow-up validation. As extensively validated in many studies, since protein activity is computed based on the coherent differential expression of a large number of transcriptional targets of regulatory proteins (regulons), VIPER-nominated MRs are expected to represent a more robust measure of the host-cell response to viral infection, compared to gene expression. The latter is often noisy and highly sparse, especially in scRNA-Seq profiles, due to gene dropout effects caused by low sequencing depth. Critically, the combination of VIPER-based MR nomination and large-scale, PLATE-Seq-based drug perturbations profiling in human airway organotypic cultures allowed identification of FDA-approved drugs representing optimal MR protein activity inverters in SARS-CoV-2 infected cells.

The population cell-specific host response signatures identified by the analysis characterize a complex interplay between innate anti-viral defense mechanisms and pro-viral host-cell mechanisms hijacked by the virus. Distinguishing these two facets remains a significant challenge due to their ambiguous dual functions (antiviral or proviral activities). Moreover, it is noteworthy that in the Reactome Pathway Enrichment analysis we found “Innate Immune System” responses significantly deactivated in the infected cells compared to bystanders. In contrast the "Cytokine Signaling in Immune System" pathway, which includes interferons and related factors, was not significantly affected, either positively or negatively, by some of the drugs identified by ViroTreat analysis (**Fig. 5d**). Indeed, top candidate MR activity inverter drugs—such as vonoprazan fumarate, aprepitant, and anisodamine hydrobromide—clearly induced inactivation of “Innate Immune System” but not “Cytokine Signaling” pathways, suggesting potential involvement of alternative mechanisms other than interferon-mediated.

ViroTreat identified 11 drugs as potentially able to rescue the normal state of regulatory of host cell programs hijacked by SARS-CoV-2 infection. Notably, four of these drugs (bedaquiline, tolperisone HCl, prasugrel, anisodamine), selected by our unbiased analysis, were also independently identified by protein-docking analyses as high-affinity binders of Mpro, the main protease of SARS-CoV-2^77^. Since our drug perturbation studies were not performed in the presence of the virus, this suggests that, in addition to interfering with Mpro—thus playing a crucial direct role in modulating viral replication—these drugs may provide a dual mechanism to attenuate COVID-19 disease severity, i.e., by both rescuing physiologic regulatory programs in infected cells and by attenuating the proteolytic activity of SARS-CoV-2. Moreover, two of the 11 drugs identified by the analysis have been clinically employed for treating SARS-CoV-2 infections, thus confirming the predictive efficacy of the ViroTreat algorithm in identifying promising candidates for therapeutic intervention against SARS-CoV-2.

Taken together, our approach using relevant primary airway culture system and network-based computational approaches to identify SARS-CoV-2-host interactions and pharmacological targets is likely to have a broader application as a platform to study the effect of various viruses in the respiratory epithelium. The value of this approach is that, rather than targeting viral proteins, which are under significant mutational stress and may adapt rapidly to avoid neutralizing antibodies or viral-protein inhibitors, drugs predicted by this approach target the cell host system to reduce both the virus ability to replicate and the mediators of the adverse epithelial response to the virus.

Finally, the gene expression profiles generated by drug perturbation of organotypic cultures of primary human airways epithelial cells represent a novel universal resource to identify candidate MR-inverter drug for virtually any airways-specific pathogen, conditional only to the availability of infected vs. non-infected cell signatures.

## METHODS

### Human airway epithelial organotypic cultures in air-liquid Interface

Primary human airway epithelial progenitors (basal cells) were isolated from de-identified adult human lung transplant organ donors in compliance with guidelines established by the International Institute for the Advancement of Medicine (IIAM). Airway progenitors from regionally distinct sites (upper, lower trachea, extrapulmonary main bronchus) were collected from different donors and tested in organotypic culture. Samples from lower trachea were used for these assays. Basal cells were cultured under submerged conditions in Pneumacult-Ex Plus expansion media (Catalog #05040). After expansion, human airway epithelial cells were dissociated with TrypLE (ThermoFisher #12604013) and plated (5 = 10^4 cells) on collagen-I-coated Transwells (0.4 micron pore, Corning #354236) and subsequently replated in 24-well Transwell plates cultured in PneumaCult Ex-Plus media under submerged conditions until confluency (day 7). Differentiation was induced by removing the medium from the top chamber and adding Pneumacult-ALI media (Catalog #05001) to the bottom chamber. Medium was changed every other day until ALI day 21, when cultures were fully differentiated, as previously described ^34–35^.

### SARS- CoV-2 Infection of airway epithelial organotypic cultures

ALI day 21 airway epithelial organotypic cultures were inoculated with SARS- CoV-2 (“Seattle strain”, NR-52281) on the apical surface at a previously tested multiplicity of infection (MOI) of 0.1. After 24 hours, the virus-containing media in the apical chamber was removed and cells were washed twice with phosphate-buffered saline (PBS) and cultured for 1, 3, and 6 dpi. Mock infection conditions at days 1, 3, and 6 were performed in parallel using the same reagents without virus. Plaque assays were performed for quantitation of viral infection as previously reported (Castaneda et al. iScience, 2023). Briefly, supernatant from the apical chamber of the Transwell culture plates was collected at the day of cell harvest and diluted to a concentration gradient at the power of 10 in 1X PBS containing 1% Bovine serum, Samples were overlaid on pre-seeded VeroE6 cell monolayers in 2% Oxoid agarose mixed with 2X MEM 0.3% FBS and incubated for 72h at 37°C. After formaldehyde fixation plaques were visualized by immunostaining with SARS-CoV-2 NP antibody (1C7). All SARS-CoV-2 infection experiments were conducted in a BSL-3 facility (Icahn School of Medicine at Mount Sinai, NY).

### Immunofluorescence

Transwell membrane inserts from human air-liquid interface organotypic cultures were fixed in 4% paraformaldehyde in PBS at room temperature for 1 hour (day 0) or overnight (day 28) and processed as reported in Zhou et al^78^. The inserts were cut into 6-8 pieces, blocked with 1% bovine serum albumin (Sigma #A3294) and 0.5% TritonX-100 (Sigma #9002-93-1) for 1 hour at room temperature. For immunofluorescence (IF) samples were incubated with anti-SARS-CoV-2 Nucleoprotein primary antibody (CTAD Mount Sinai #NP-1C7) in 1% bovine serum albumin (Sigma) and 0.5% TritonX-100 at 4°C overnight, then washed with PBS and incubated with Alexa Fluor-conjugated secondary antibodies (1:300) and NucBlue Live Cell ReadyProbes Reagent (DAPI) (Life Technology) for 1 hour. After washing, samples were mounted with ProLong Gold antifade reagent (Life Technology).

### Sample preparation for scRNA-Seq

The apical chamber of Transwell plates containing either Infected or mock-infected cells were washed 2x with PBS at 37C for 5 mins. PBS was then removed and 500 ul of accutase containing 5 mM EDTA was added into the apical and basolateral sides of the Transwells. The cells were gently pipetted up and down, collected into a 15 ml conical tube, and centrifuged at 1500 rpm for 3 minutes at 4 ^0^C. The medium was aspirated, and cells were resuspended in 500 ul DMEM+10% FBS and filtered through a 04 ul cell strainer into a 1.5 mL. tube. Cells were then counted and diluted to 1 million cells/mL and placed on ice and sent for single cell RNA seq (10X Genomics).

### scRNA-Seq pre-processing and annotation of cell types and infection status

scRNA-Seq FASTQ files were aligned to the human reference genome (GRCh38-2020-A) using 10x Genomics CellRanger software (v 5.0.1) to generate raw count matrices for all the four different experimental conditions, MOCK, 1dpi, 3dpi and 6 dpi. Before proceeding with downstream analyses, stringent quality-control (QC) filtering metrics were applied to the data to remove low quality cells. Specifically, cells with less than 30% content of mitochondrial genes and more than 1,000 UMIs were retained. After QC filtering the average number of unique molecular reads per cell (UMIs/cell) was 2,723.742 and the average number of detected genes per cell was 1,053.218. Cells were labelled as either a basal, secretory or ciliated cell according to the results of Gene Set Enrichment Analysis (GSEA) with manually curated well-established marker genes for each epithelial cell subtypes. As independent confirmation of the cell types inferred using GSEA, we have also trained SingleR to annotate our single-cell data from MOCK condition using reference signatures published in Ravindra et al.^34^.

To quantify the number of infected cells in each experimental condition, reads in the FASTQ files were re-mapped to the SARS-CoV-2 reference genome (NC_045512.2) from GeneBank^79^. For this, we used Kallisto Bustools^80^ to estimate gene expression at the single-cell level. Infected cells were initially identified by detecting at least one read in any SARS-CoV-2 gene. Also, we analyzed the proportion of infected cells across the three cell types as viral read thresholds increased. At each threshold, a consistent proportion of infected cells was observed.

### Airway network inference from microarrays and scRNA-seq data

We generated gene regulatory networks specific for human airways using an ARACNe algorithm with Adaptive Partitioning^81^ (ARACNe-AP). In brief, ARACNe-AP determines protein-gene regulatory interactions by assessing the statistical significance of mutual information (MI) between the gene expressions of regulator proteins and potential targets. Subsequently, the algorithm eliminates statistically significant candidate targets that violate the data processing inequality. For accurate mutual information analysis by the ARACNe-AP algorithm, a minimum of 100 expression profiles is typically required. The output of ARACNe-AP consists of the likelihood and the regulatory action mode of each protein-gene interaction. The likelihood is an edge weight, ranging between 0 and 1, corresponding to the scaled MI score that is divided by the maximum MI in all edges. The regulatory action mode is the sign of the association (>0: induction, <0: inhibition) between the protein regulator and its target gene, ranging between -1 and +1, computed by spearman correlation. For an input of ARANCe-AP, we retrieved publicly available gene expression data of human bronchial and nasal epithelial samples from two studies illuminating diagnostic markers of lung cancer^37,38^ (GEO accession numbers: GSE66499 and GSE80796). The downloaded data had been normalized using robust multiarray analysis^82^ (RMA) and scaled into log_2_ intensities. Among the samples, we inferred the networks, only utilizing benign lung disease samples (*n* = 190 for GSE66499, *n* = 196 for GSE80796) in each data set, separately. Also, we prepared an input of the regulator protein list, consisting of the transcriptional (co)regulaors identified based on the following Gene Ontology (GO) terms: transcription regulator activity (GO:0140110), transcription coregulator activity (GO:0003712), and DNA-binding transcription activator activity (GO:0001216). ARACNe-AP was implemented with the setting of 200 bootstraps and Bonferroni-corrected threshold of *p* = 0.05 for detecting statistically significant protein-target interactions. Then, we merged the regulatory networks inferred from two datasets (GSE66499 and GSE80796) by averaging the estimates, namely likelihoods and regulatory action modes for each protein-gene interaction across the two networks. To ensure accuracy, we limit regulons, a set of target genes for each regulator, to a maximum of 50 genes, as the improvement in the precision of VIPER analyses plateaus beyond this point. In addition, we only considered the edges with a strong weight (i.e. likelihood>0.25). Consequently, only the 50 most statistically significant targets within each regulon were retained. This cautious approach prevents potential biases in VIPER NES assessment, as NES measured from larger regulons would exhibit higher statistical significance.

Similarly, we reverse-engineered gene regulatory networks from scRNA-Seq data across four conditions (MOCK, 1dpi, 3dpi, and 6dpi), using ARACNe-AP. To address the sparsity in single-cell expression, we first created metaCells by merging 10 neighboring cells per condition. ARACNe-AP was then applied to the metaCell expressions, producing a network for each of the four conditions, which were subsequently used in metaVIPER analysis to infer single-cell protein activity profiles, as shown in **Suppl . Fig.2a**.

### Generation of Host Response SARS-CoV-2 Signatures

First, we generated differential gene expression signatures by comparing the expression profiles of infected and non-infected cells in each cell type. Cell type-specific clusters were identified by applying cell marker enrichment analysis to Louvain clustering of protein activity profiles, inferred through metaVIPER with scRNA-based networks as previously described.

A ‘Bootstrapped Approach’ was implemented in which we compared *k* randomly selected infected cells at each time point with the non-infected cells nearest to each infected cell, using a two-sample Mann-Whitney U test, repeated 100 times. In other words, we generated 100 metaCell pairs, each comparing the mRNA reads of *k* random sampled infected cells and of *k* non-infected cells from each time point, using the Mann Whitney U-test (**Fig. 2A**).

For the infected-vs-mock signature, the non-infected cells were sampled from the mock condition, based on the distance in the PCA space and the single-cell VIPER-inferred protein activity. For the infected-vs-bystander signature, the uninfected bystander cells nearest to the infected cells were sampled from the cells with no viral read detection at the same time point.

To determine an optimal *k*, we iterated the above procedure with increasing *k* one-by-one. As *k* increases, the differential gene expression converges, and *k* = 23 was determined as an optimal number. In other words, the optimal value of *k* (*k*_Opt_ = 23) was determined by assessing convergence of the metaCell-based analysis vs. analyzing the differential expression of all infected vs. all non-infected cells. We integrated the bootstrapped signatures using the Stouffer’s method for each combination of cell type and timepoint. Differential protein activity was then integrated across all 100 metaCells using Stouffer’s z-score integration method. These signatures were then used for further analyses.

### Hallmark pathway enrichment analysis

To gain insights into the biological pathways that are activated and deactivated during SARS-CoV-2 infection, we conducted pathway enrichment analysis using hallmark gene sets obtained from Molecular Signatures Database (MSigDB; https://www.gsea-msigdb.org/)^83^. NES of hallmark pathways in each host response signature were computed using *analytic Rank-based Enrichment Analysis (aREA)*^10^ in the VIPER R package. The signs and weights for individual genes for each hallmark set were set to ones during the *aREA* implementation.

### Enrichment analysis of proviral genes

We compiled a set of proviral host factors from CRISPR-KO assays previously conducted on human lung cancer cell lines (A549 and Calu-3) ^3,31,32^. Specifically, we identified genes with a z-score >1.5, reported in at least one of three studies (Rebendenne et al.^3^, Daniloski et al.^32^, Biering et al.^31^) as proviral genes, where the z-score reflects gene essentiality during SARS-CoV-2 infection. Among these host factors, 50 transcription factors (TFs) and cofactors were identified. We then conducted enrichment analysis of the 50 TFs and cofactors using the fast GSEA (fgsea)^84^ R package, visualizing the results through enrichment plots (the *plotEnrichment* function). Additionally, leading-edge proteins were identified as those ranked highest up to the point of maximum enrichment score among the 50 TFs and cofactors.

### Drug perturbation screen of airway organotypic cultures and PLATE-seq data pre- processing

Drug perturbations were performed in fully differentiated ALI day 21 organotypic airway and analyzed 24 hours later. C_max_ represents the maximum tolerated serum concentration of each drug as determined from established studies. Drug signatures were generated using PLATE-Seq, a high-throughput RNA-Seq platform that allows simultaneously profiling multiple drugs^74^.A total of 441 drugs, 20 DMSO, and 20 untreated samples were sequenced over ten 96-well plates with duplicates. Note that drugs in each plate were randomly selected regardless of their mechanism of action.. Since samples in 96 wells of each plate were pooled together in PLATE-Seq technology, we demultiplexed the reads using the computational tool Sabre (https://github.com/najoshi/sabre/) using the sample barcode identifiers. Then, we applied kallisto^85^ to quantify read counts and gene expression abundance matrices (e.g. transcripts per million; TPM) for each plate. In detail, while running kallisto, we first generated the human transcriptome indices from the human genome reference (*Homo_sapiens.GRCh38.cdna.all.fa* and *Homo_sapiens.GRCh38.96.gtf* as of Mar/12/2019) in Ensembl^86^, using *kallisto index*. The transcriptome indices were used during *kallisto quant* to quantify pair-end reads of the PLATE-seq data.

### Protein activity inference for PLATE-seq data

We inferred protein activity profiles from the PLATE-seq data, using VIPER^10^. VIPER computes the normalized, rank-based enrichment score (NES) of the regulon of each protein in genes differentially expressed when comparing treatment versus a control. Statistically significant positive and negative NES values provided by VIPER imply activated or inactivated proteins in the state of interest compared to the control state. Meanwhile, non-significant NES scores signify proteins with no significant change in activity. Unlike a conventional GSEA, VIPER employs a probabilistic model to integrate consensus among activated, inhibited, and unclearly regulated targets, considering their differential expression. As a result, utilizing a context-specific network which can enhance interactome fidelity is critical for accurately inferring protein activity. In this work, we utilized the airway regulatory network, inferred by ARACNe-AP as described above, for identifying protein activity profiles of PLATE-seq data. As part of the VIPER analysis, we computed the differential gene expression profiles resulting from drug treatment compared to the control state (DMSO). To mitigate the impact of technical batch effects on differential gene expression, samples were compared with DMSO conditions within the same plate. The differential expression of gene *i* for sample *j* (DE*_ij_*) was determined using the provided equation.

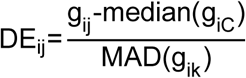

where *g_ij_* is the TPM-normalized expression of gene *i* for a treatment sample *j*, *g_iC_* is the TPM-normalized expression for gene *i* of the DMSO control samples *C*, and MAD stands for median absolute deviation (=median(|*g_ik_*-median(*g_iC_*)| and *k* ∈ *C*). To minimize denominator noise arising from low expression variance, we replaced instances where MAD(*g_ij_*) <0.01 with 0.01. Protein activity inference of the PLATE-seq data were computed using the human airway network, in the same manner as done above. For duplicated samples, we averaged the protein activity between them.

### ViroTarget

For the viral-induced host response signature, druggable MRs were identified based on the targets of 1,738 known drug inhibitors from DrugBank^69^. To identify inhibitable MRs in our signature, we excluded those with absent gene expression or negative protein activity. Consequently, the final selection of druggable MRs comprised targets significantly activated (Benjamini-Hochberg corrected p-value < 0.05) and expressed (read count > 0) in at least one cell type among ciliated, basal, and secretory cells.

### ViroTreat

To predict drugs capable of reversing the host response signature to SARS-CoV-2 infection, we employed ViroTreat, a modified version of the OncoTreat pipeline^18^. The pipeline fundamentally utilizes two data inputs: (1) a target signature and (2) drug perturbation signatures. In our study, the target signature comprised the top 25 activated and 25 inactivated MRs derived from the host response scRNA-Seq data. Simultaneously, the drug signatures were constructed based on protein activity profiles of all transcriptional regulators using PLATE-seq data. The statistical score of the host response inversion against individual drug signatures was computed using *aREA*. The NES serves as a measure of similarity, with a more negative value indicating a stronger inversion of the host response signature relative to the corresponding drug signature. This computation utilized the *aREA* function, a weighted GSEA algorithm in the VIPER R package. In the *aREA* implementation, we assigned a value of +1 to the top 25 activated MRs and -1 to the top 25 inactivated MRs in the host response signature. We also applied a penalty to MRs whose activities were consistently activated or deactivated with any drug treatment (i.e. low specificity), reducing their importance by adjusting their weight through a sigmoid function during the *aREA* application. Finally, we predicted drugs through this pipeline, separately to the host response from each cell type: basal, secretory, and ciliated cells.

### Reactome pathway enrichment analysis

To identify pathways significantly modulated by the 11 drugs which were identified as strong host-response inverters by ViroTreat, we computed NES of reactome pathway^76^ terms in each drug signature. For this, we used the *gsePathway* function in the ReactomePA^87^ R package. Prior to implementation, the drug signature was sorted in descending order of protein activity.

## Data Availability Statement

All data generated for this manuscript is available upon request to the corresponding authors.

## ACKNOWLEDGEMENTS

NIH-NHLBI R35-HL135834 to W.V.C and NIH-NCI R35 CA197745, S10 OD012351,S10 OD021764 to A.C. MS and AGS want to thank Randy Albrecht for management and organization of the BSL3 facility. SARS-CoV-2 work in the M.S. laboratory is supported by NIH/NIAID R01AI160706, NIH/NIDDK R01DK130425 and U19AI135972 to AGS.

## AUTHOR CONTRIBUTIONS

WVC, AC, AGS conceived and designed the study. BD, DC, YZ, JQ, XK generated primary airway organotypic cultures. HN, LT perfomed computational and statistical analyses. SJ and MS performed the SARS-CoV-2 infection and viral titration of organotypic cultures. JQ, DC performed histological and IF analysis. BD, CK performed plate seq and drug perturbation analyses. BD, HN, WVC, AC interpreted results. BD, HN, LT generated Figures. BD,HN, LT, AC, WVC co-wrote the manuscript.

---

**Supplementary Figure 1: Adult human airway epithelial cell populations in control and SARS-CoV-2-exposed ALI cultures**

A. Diagram: airway epithelial cell types analyzed in the ALI organotypic cultures and panel of markers used for their identification.

B. Top panels: PCA plots of GSEA results using the gene expression signature in each cell type; high (red) and low (blue) scores are depicted. Lower panels: feature plots of representative marker genes with expression in transcripts per million represented in a gradient from red (high) to gray (low).

C. PCA plot of the independent validation of Cell Type Annotation using an external Single-Cell dataset (Ravindra et al.^34^) as reference to train SingleR^33^.

D. Plot showing the percentage of infected cells (y-axis) in the three different cell types (basal ciliated, secretory) when the threshold of viral reads ranges from 1 to 100 (x-axis) at either 3 or 6 dpi.

**Supplementary Figure 2: Analyses of SARS-CoV-2 host responses in airway organotypic cultures and other cell types.**

A. PCA plots based on protein activity of quality-control filtered cells to demonstrate consistency between protein activity and gene expression. **Left panel:** cells colored according to the timepoints (MOCK, 1 dpi, 3 dpi, 6 dpi). **Middle panel**: cells colored based on cell types: Basal, Ciliated, and Secretory (>75% of the cells showing agreement between gene expression and protein activity. **Bottom panel**: PCA plot based on protein activity data of infected (Red) and non-infected (Grey) cells.

B. **Top panel**: Scatterplot of hallmark-normalized enrichment score comparisons based on NES protein activity (x-axis) and gene expression analysis (y-axis) in SARS-CoV-2 *treated and mock cells* at 3 and 6dpi. IFN alpha and IFN gamma response pathways were consistently enriched in all cell types by both gene and protein activity. **Bottom panel**: Scatterplot of hallmark-normalized enrichment score comparisons based on NES protein activity (x-axis) and gene expression analysis (y-axis) in SARS-CoV-2 *infected vs uninfected* cells at 3dpi. R and p-values denote a spearman correlation coefficient and its statistical significance, computed using the *stats*^88^ package in R.

C. Heatmap showing a comparison of the host response signatures (top 25 activated and top 25 inactivated MRs) of SARS-CoV-2 *infected vs. mock* across different airway epithelial cell types in ALI cultures (3 and 6dpi), human airway-derived lung cancer cells (calu-3) and gastrointestinal organoids (ileum, colon) using aREA analysis (see Methods). Positive (green) and negative (purple) NES values are indicated in heatmap. The asterisk symbols denote the following p-value thresholds: *: p<0.05, **: p<0.01, and ***: p<0.001.

**Supplementary Figure 3: Proviral host factors are induced in overlapping and distinct airway epithelial cell types**

Venn-diagrams displaying proviral factors enriched in the leading edge of the host response signature of SARS-CoV-2 *infected vs. bystander* (blue) or SARS-CoV-2 *infected vs. mock* (yellow) in each cell type at 3 dpi. The percentage of proviral factors identified by both approaches (intersection) in each cell type is indicated in ciliated (29.4%), secretory (23.5%) and basal (59.0%) cells.

**Supplementary Figure 4: ViroTarget analysis of druggable MRs of the SARS-CoV-2 host responses.**

Heatmap summarizing the druggable MR candidates identified by the ViroTarget algorithm in the SARS-CoV-2 *infected vs. bystander* cells in each cell type (see methods). The heatmap includes druggable MRs enriched in at least one signature of host responses in ciliated, basal, and secretory cells. Their activity is shown in red (activated) and blue (deactivated) across cell types. Candidate drugs identified as an inverter of the activity of each MR are shown on the table below.

**Supplementary Figure 5: ViroTreat analysis of druggable MRs of SARS-CoV-2 host responses.**

A. ViroTreat enrichment plots for the 11 drugs identified as significant inverters of the host SARS-CoV-2 host response in *infected vs. bystander* (SIS) in each of the cell types indicated

B. Heatmap of candidate drugs predicted by ViroTreat as inverters of the host SARS-CoV- 2 host response *in infected vs. bystander* (SIS) cells at 3 dpi in each cell type. Asterisks indicate *p<0.05, **p<0.01, ***p<0.001.

**Supplementary Figure 6: ViroTreat analysis of drugs reversing MRs and proviral factors of the SARS-CoV-2 host responses.**

A. Heatmap of the VIPER-inferred top 25 differentially activated (left) and inactivated (right) MR proteins of SARS-CoV-2 *infected vs. bystander* cultures (SIS) at 3 dpi and ViroTreat- inferred top 11 drugs predicted to revert the activity of these MR in each cell type (basal, secretory, ciliated). The MR-reversal scores are represented in the grid if activated (red) or inactivated (blue) by each of the 11 drugs. Numbers in the boxes refer to the rank of the protein in the inverted signature. Reversal of the MRs was statistically significant (p< 0.05 Benjamini-Hochberg test).

B. Heatmap of the protein activity of proviral factors found in the leading edge of SARS-CoV- 2 *infected vs. bystander* for each cell type at 3dpi (left). MR-reversal of these proviral factors by the top 11 drugs predicted by Virotreat, represented in the grid as above (right). Reversal of the MRs was statistically significant (p< 0.05 Benjamini-Hochberg test).

