## Supplemental Figures and Tables for "Identification and Targeting of Regulators of SARS-CoV-2-Host Interactions in the Airway Epithelium"

a

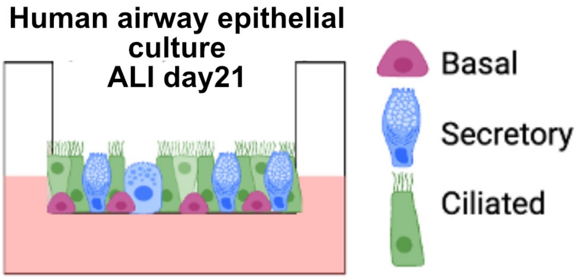


cell-type markers

| Basal | Secretory | Ciliated |
|---|---|---|
| TP63 | MUC5AC | FOXJ1 |
| KRT5 | MUC5B | TUBB4B |
| ITGA6 | GP2 | TUBA1A |
| NGFR | SPDEF | CDHR3 |
| PDPN | TFF1 | PROM1 |
| KRT17 | SCGB1A1 | |
| KRT14 | SCGB3A2 | |
| | CYP2F2 | |

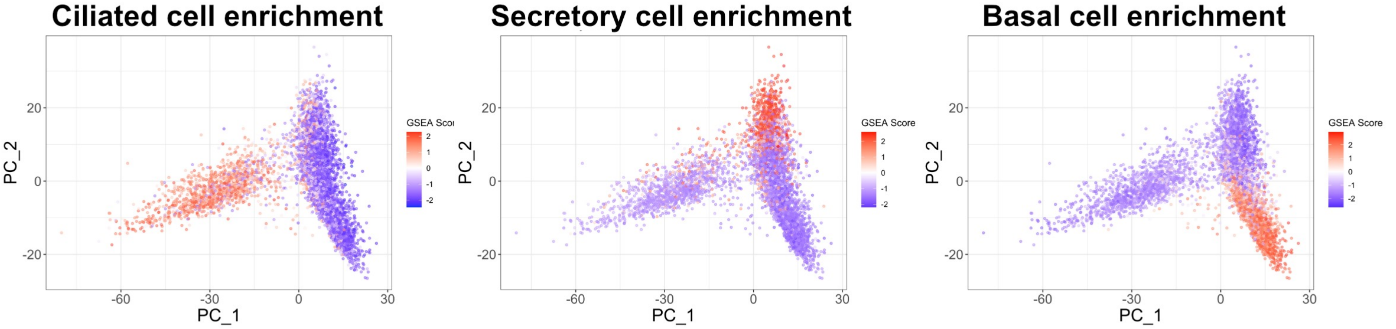


b

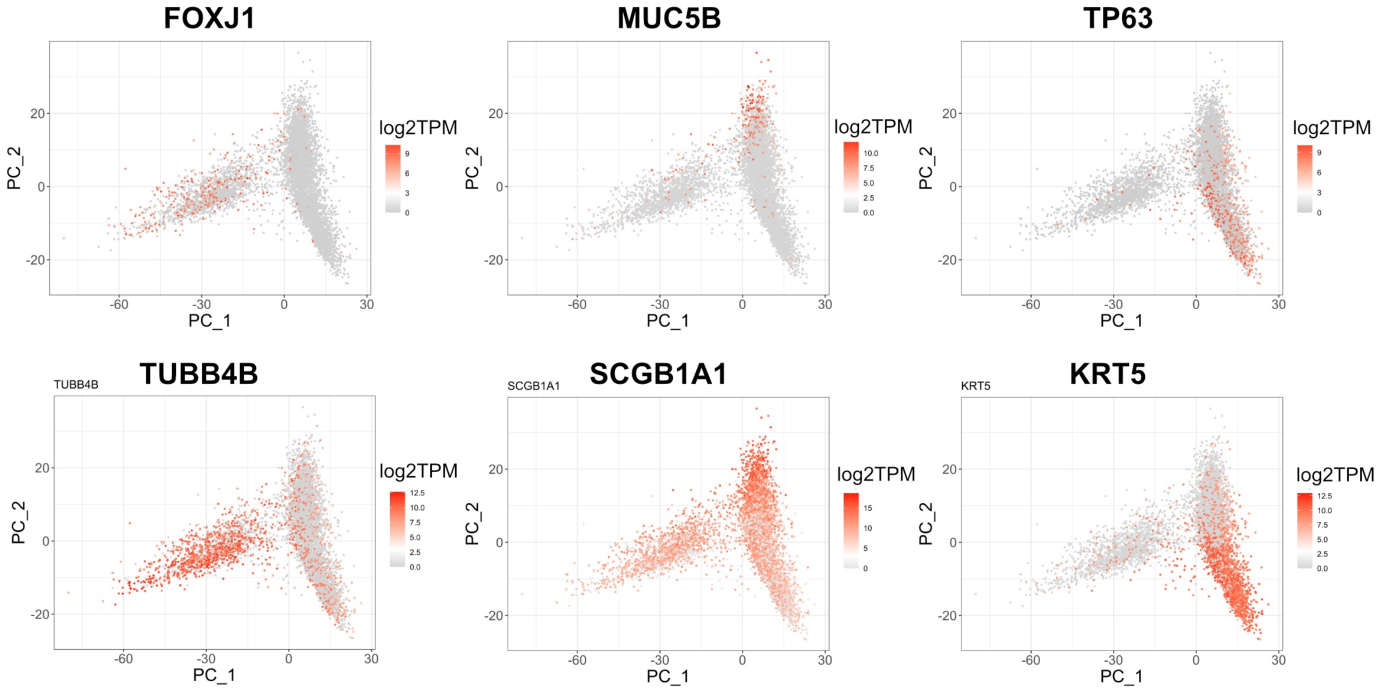


c

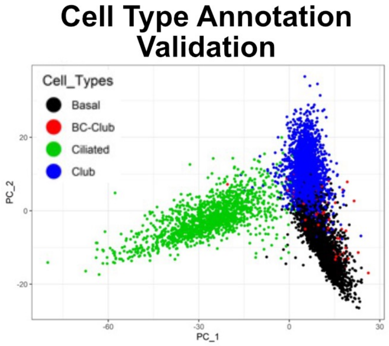


D

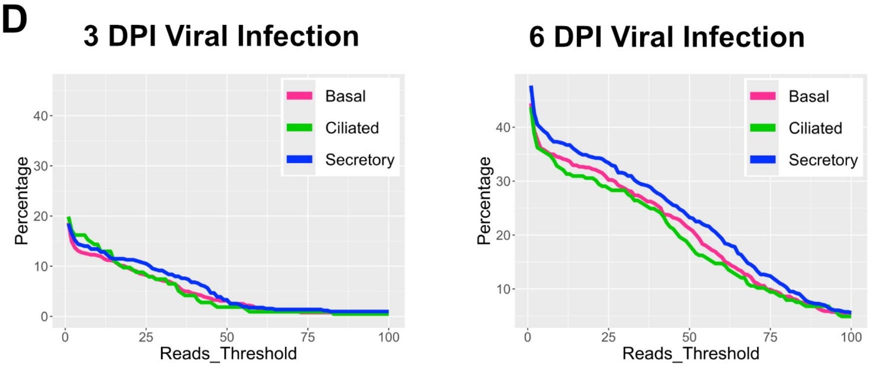


### Supplementary Figure 1

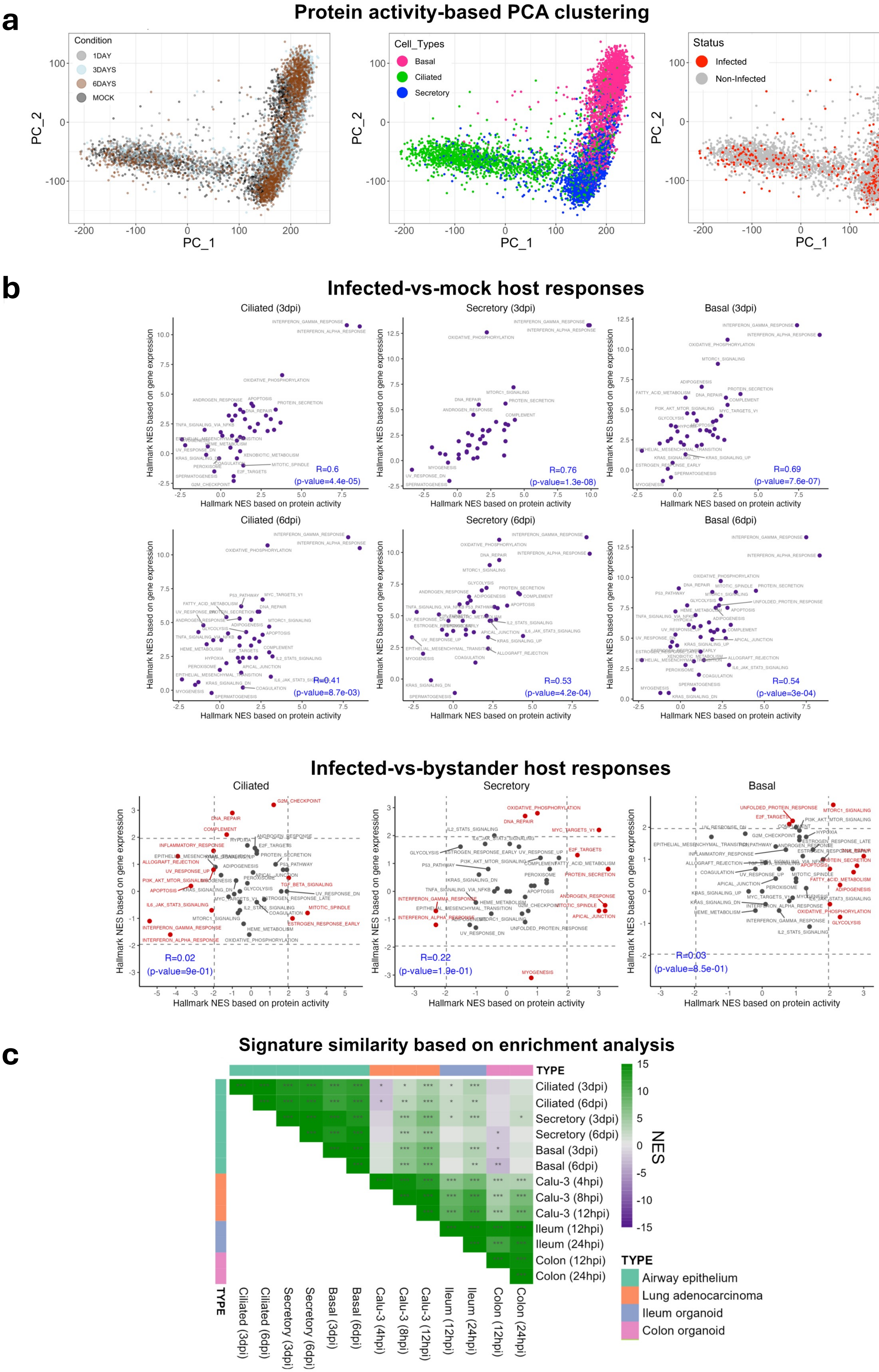


### Supplementary Figure 2

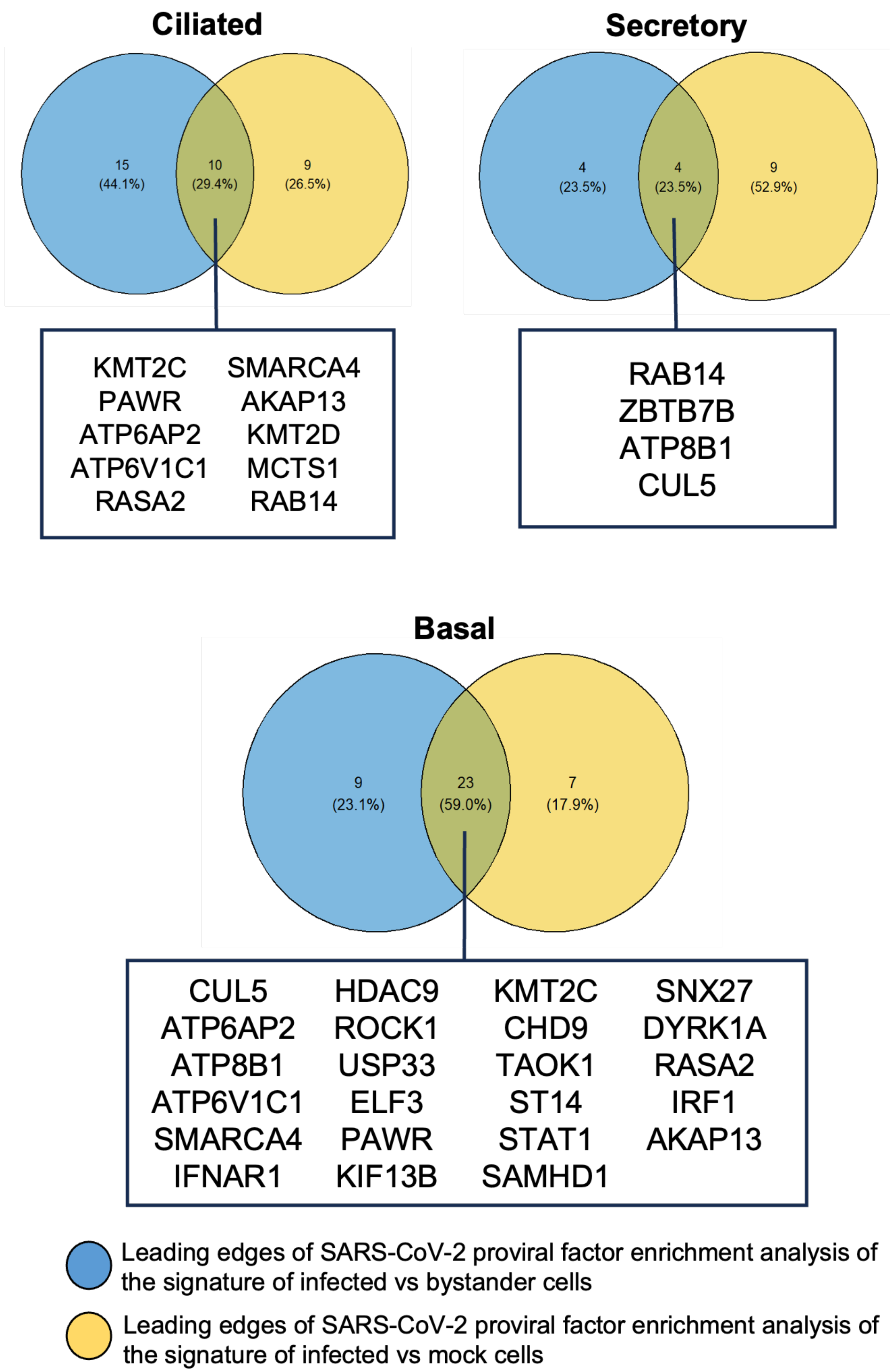


**Supplementary Figure 3**

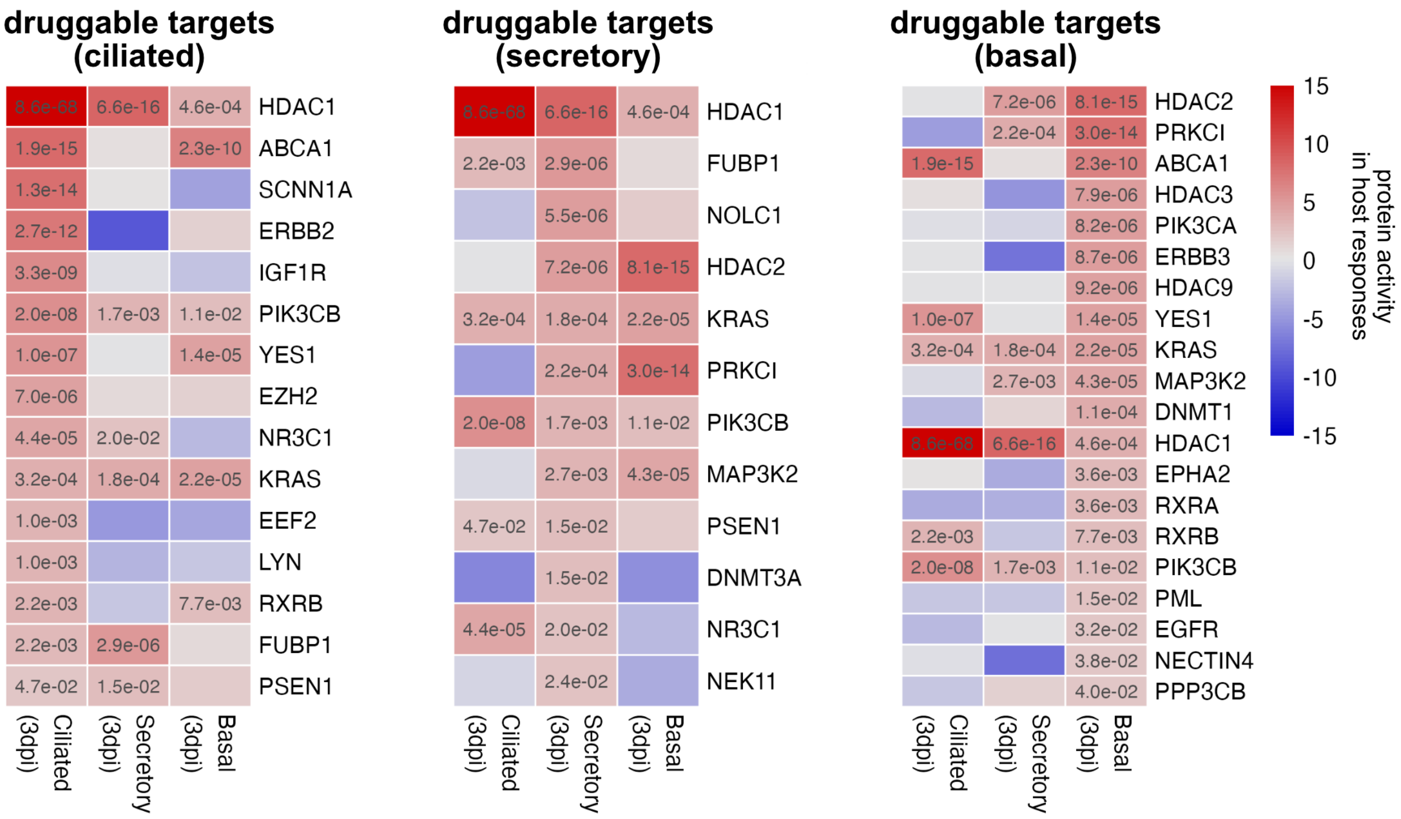


| druggable taget | inhibitors |
|---|---|
| ABCA1 | Probucol |
| DNMT1 | Azacitidine, Decitabine, Thioguanine |
| DNMT3A | Azacitidine, Decitabine |
| EEF2 | Moxetumomab pasudotox |
| EGFR | Cetuximab, Erlotinib, Panitumumab, Afatinib, Dacomitinib, Osimertinib, Necitumumab, Brigatinib, Mobocertinib, Amivantamab, Gefitinib, Lapatinib, Vandetanib, Neratinib |
| EPHA2 | Dasatinib, Regorafenib |
| ERBB2 | Trastuzumab, Ado-Trastuzumab emtansine, Margetuximab, Tucatinib, Lapatinib, Pertuzumab, Afatinib, Neratinib, Dacomitinib, Osimertinib, Brigatinib |
| ERBB3 | Osimertinib |
| EZH2 | Tazemetostat |
| FUBP1 | Sacituzumab govitecan |
| HDAC1 | Belinostat, Panobinostat, Romidepsin, Vorinostat, Decitabine |
| HDAC2 | Belinostat, Panobinostat, Romidepsin, Vorinostat, Valproic Acid, Decitabine |
| HDAC3 | Belinostat, Panobinostat, Vorinostat, Romidepsin, Decitabine |
| HDAC9 | Valproic Acid, Belinostat, Panobinostat, Romidepsin, Decitabine, Vorinostat |
| IGF1R | Brigatinib |
| KRAS | Sotorasib, Adagrasib |
| LYN | Dasatinib, Bosutinib, Ponatinib, Nintedanib |
| MAP3K2 | Bosutinib |
| NECTIN4 | Enfortumab vedotin |
| NEK11 | Dabrafenib |
| NOLC1 | Doxorubicin |
| NR3C1 | Mifepristone |
| PIK3CA | Alpelisib, Copanlisib, Idelalisib, Quercetin |
| PIK3CB | Copanlisib, Idelalisib, Quercetin |
| PML | Arsenic trioxide |
| PPP3CB | Pimecrolimus |
| PRKCI | Tamoxifen |
| PSEN1 | Nirogacestat |
| RXRA | Bexarotene |
| RXRB | Bexarotene |
| SCNN1A | Triamterene, Amiloride |
| YES1 | Dasatinib |

### Supplementary Figure 4

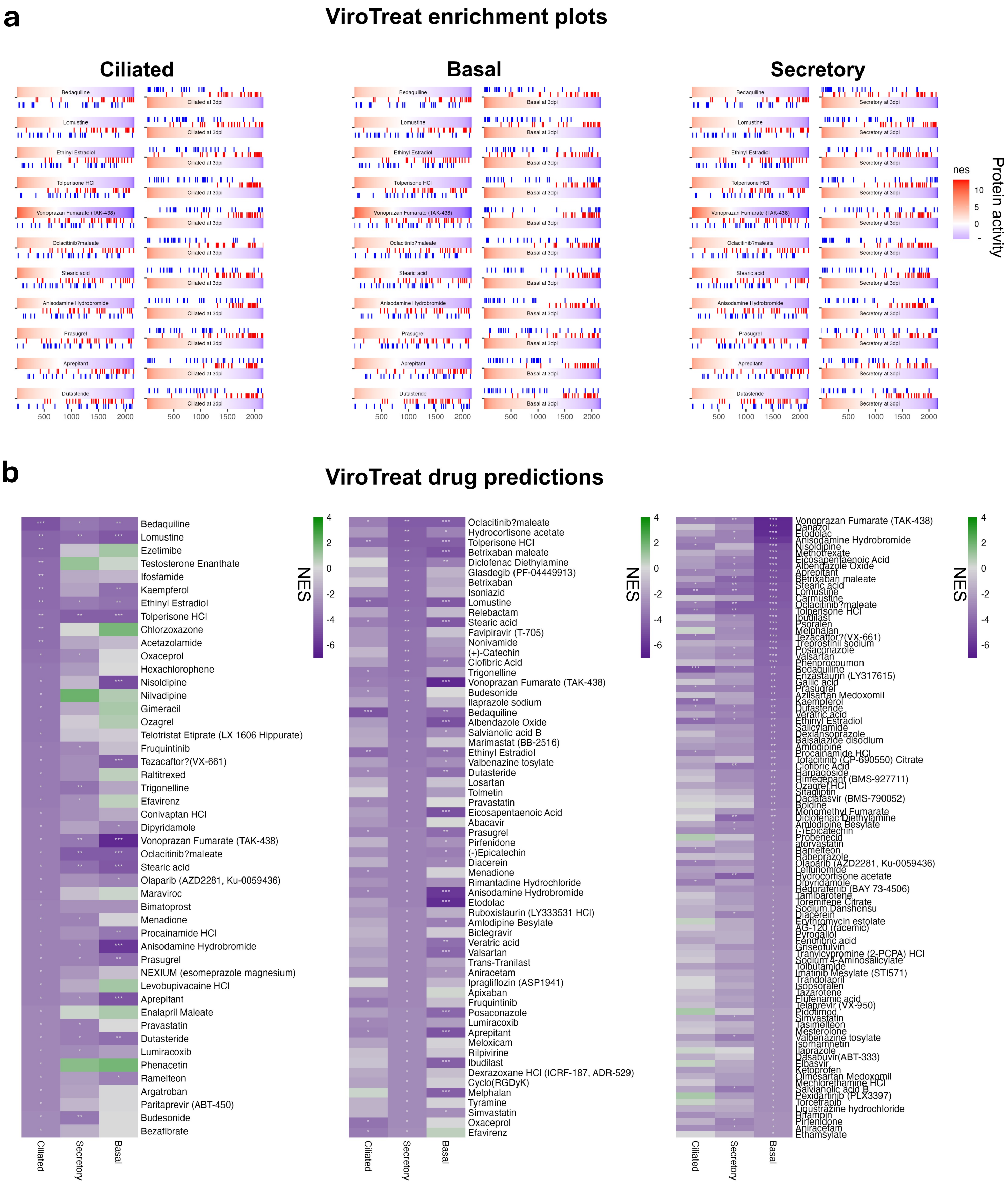


### Supplementary Figure 5

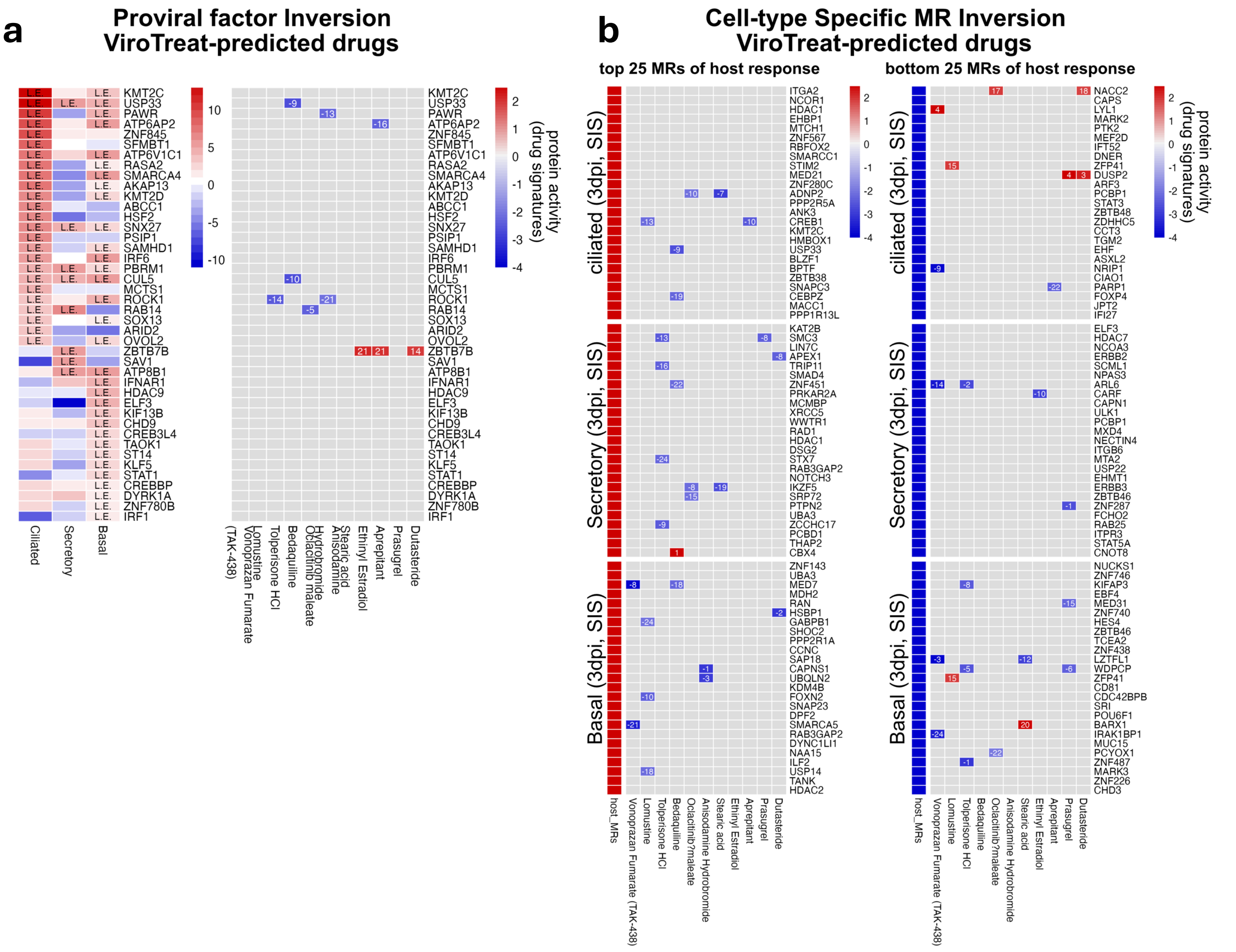


### Supplementary Figure 6

### Supplementary Table 1

| Condition | Quality Control Status | Number of Cells | Mean #UMIs/cell | Mean #Detected Genes/cell |
|---|---|---|---|---|
| **MOCK** | Pre-QC | 2545 | 1850.103 | 824.0998 |
| | **Post-QC** | **1769** | **2293.083** | **974.9638** |
| **1 DPI** | Pre-QC | 2512 | 1682.525 | 774.541 |
| | **Post-QC** | **1628** | **2180.012** | **938.6229** |
| **3 DPI** | Pre-QC | 1895 | 2131.499 | 880.381 |
| | **Post-QC** | **1373** | **2624.546** | **1034.783** |
| **6 DPI** | Pre-QC | 2396 | 2947.942 | 1036.608 |
| | **Post-QC** | **1625** | **3821.113** | **1268.788** |
| **All Conditions** | Pre-QC | 9348 | 2143.504 | 876.6597 |
| | **Post-QC** | **6395** | **2723.742** | **1053.218** |

### Supplementary Table 2

| | MOCK | 1 DPI | 3 DPI (total) | 3 DPI (Infected) | 6 DPI (total) | 6 DPI (infected) |
|---|---|---|---|---|---|---|
| **Basal** | 535 | 405 | 643 | 120 (18.66%) | 698 | 310 (44.41%) |
| **Ciliated** | 503 | 523 | 216 | 43 (19.9%) | 265 | 116 (43.77%) |
| **Secretory** | 731 | 700 | 514 | 95 (18.48%) | 662 | 316 (47.73%) |

### Supplementary Table 3a

#### Top Activated and inactivated MRs in the host response signatures **Infected vs Mock**

**Top 25 activated proteins**

| | Ciliated (3dpi) | | | Ciliated (6dpi) | | | Secretory (3dpi) | | | Secretory (6dpi) | | | Basal (3dpi) | | | Basal (6dpi) | | |
|---|---|---|---|---|---|---|---|---|---|---|---|---|---|---|---|---|---|---|
| rank | protein | NES | FDR | protein | NES | FDR | protein | NES | FDR | protein | NES | FDR | protein | NES | FDR | protein | NES | FDR |
| 1 | IFITM1 | 4.67E+01 | 0.00E+00 | BST2 | 2.34E+01 | 1.43E-118 | IFITM1 | 4.87E+01 | 0.00E+00 | MX1 | 2.29E+01 | 2.73E-112 | IFI6 | 5.28E+01 | 0.00E+00 | ZNFX1 | 3.52E+01 | 1.23E-268 |
| 2 | IFI6 | 4.67E+01 | 0.00E+00 | IFI27 | 2.27E+01 | 1.00E-111 | IFI6 | 4.76E+01 | 0.00E+00 | ZNFX1 | 2.20E+01 | 6.33E-104 | MX1 | 5.00E+01 | 0.00E+00 | IFI27 | 3.33E+01 | 1.17E-240 |
| 3 | IFI27 | 4.62E+01 | 0.00E+00 | SP110 | 2.26E+01 | 4.18E-111 | MX1 | 4.65E+01 | 0.00E+00 | IFI6 | 2.18E+01 | 2.42E-102 | IFI27 | 4.88E+01 | 0.00E+00 | IFITM1 | 3.29E+01 | 5.56E-235 |
| 4 | MX1 | 4.50E+01 | 0.00E+00 | IFITM1 | 2.09E+01 | 4.02E-95 | ZNFX1 | 4.63E+01 | 0.00E+00 | SP110 | 2.11E+01 | 9.41E-96 | ZNFX1 | 4.82E+01 | 0.00E+00 | SP110 | 3.20E+01 | 2.32E-221 |
| 5 | IRF9 | 4.48E+01 | 0.00E+00 | ZNFX1 | 2.09E+01 | 9.51E-95 | IFI27 | 4.54E+01 | 0.00E+00 | IFITM1 | 2.10E+01 | 7.29E-95 | SP110 | 4.60E+01 | 0.00E+00 | IFI6 | 3.15E+01 | 1.75E-215 |
| 6 | ZNFX1 | 4.31E+01 | 0.00E+00 | IFI6 | 2.07E+01 | 8.11E-93 | SP110 | 4.29E+01 | 0.00E+00 | OAS3 | 2.05E+01 | 1.57E-90 | IFITM1 | 4.46E+01 | 0.00E+00 | MX1 | 3.14E+01 | 3.17E-214 |
| 7 | PLSCR1 | 4.02E+01 | 0.00E+00 | IRF9 | 2.03E+01 | 4.59E-90 | IRF9 | 4.27E+01 | 0.00E+00 | IFI27 | 2.04E+01 | 7.28E-90 | STAT2 | 4.21E+01 | 0.00E+00 | PLSCR1 | 3.13E+01 | 1.10E-212 |
| 8 | SP110 | 3.96E+01 | 0.00E+00 | PLSCR1 | 2.03E+01 | 1.76E-89 | OAS3 | 4.07E+01 | 0.00E+00 | PLSCR1 | 1.94E+01 | 4.28E-81 | IRF9 | 4.01E+01 | 0.00E+00 | IRF9 | 3.06E+01 | 1.27E-202 |
| 9 | PARP14 | 3.91E+01 | 0.00E+00 | CASP7 | 2.02E+01 | 6.33E-89 | STAT2 | 3.95E+01 | 0.00E+00 | IRF9 | 1.93E+01 | 5.35E-81 | OAS3 | 3.97E+01 | 0.00E+00 | PARP9 | 2.97E+01 | 6.94E-192 |
| 10 | OAS3 | 3.80E+01 | 0.00E+00 | SP100 | 2.01E+01 | 5.54E-88 | PLSCR1 | 3.95E+01 | 0.00E+00 | STAT2 | 1.92E+01 | 4.94E-80 | PLSCR1 | 3.78E+01 | 0.00E+00 | PARP14 | 2.96E+01 | 1.86E-190 |
| 11 | STAT2 | 3.71E+01 | 5.05E-299 | TNFSF10 | 2.00E+01 | 5.49E-87 | PARP9 | 3.85E+01 | 0.00E+00 | BST2 | 1.81E+01 | 8.25E-71 | EIF2AK2 | 3.74E+01 | 2.11E-304 | SP100 | 2.91E+01 | 2.68E-184 |
| 12 | SP100 | 3.71E+01 | 5.31E-299 | STAT2 | 2.00E+01 | 8.19E-87 | PARP14 | 3.69E+01 | 1.45E-296 | PARP9 | 1.77E+01 | 1.18E-67 | PARP14 | 3.69E+01 | 4.05E-296 | OAS3 | 2.89E+01 | 4.53E-181 |
| 13 | TRIM22 | 3.64E+01 | 2.70E-287 | TRIM22 | 1.99E+01 | 3.22E-86 | BST2 | 3.63E+01 | 7.04E-286 | CASP7 | 1.69E+01 | 9.67E-62 | SP100 | 3.66E+01 | 1.37E-291 | STAT2 | 2.88E+01 | 2.41E-180 |
| 14 | PARP9 | 3.60E+01 | 2.03E-281 | KLF13 | 1.91E+01 | 2.39E-79 | CASP7 | 3.59E+01 | 1.12E-279 | TRIM22 | 1.66E+01 | 9.63E-60 | PARP9 | 3.60E+01 | 5.02E-281 | TRIM22 | 2.77E+01 | 1.09E-166 |
| 15 | BST2 | 3.55E+01 | 3.59E-274 | PARP14 | 1.90E+01 | 3.35E-79 | TRIM22 | 3.51E+01 | 1.75E-267 | SP100 | 1.65E+01 | 1.13E-58 | TRIM22 | 3.53E+01 | 3.88E-270 | TNFSF10 | 2.61E+01 | 5.20E-148 |
| 16 | EIF2AK2 | 3.18E+01 | 3.13E-219 | MX1 | 1.88E+01 | 4.81E-77 | IFITM3 | 3.29E+01 | 1.24E-235 | PARP14 | 1.60E+01 | 9.69E-56 | TNFSF10 | 3.31E+01 | 8.92E-239 | BST2 | 2.56E+01 | 5.94E-142 |
| 17 | STAT1 | 3.14E+01 | 1.93E-214 | OAS3 | 1.82E+01 | 1.14E-72 | SP100 | 3.24E+01 | 2.07E-228 | IFITM3 | 1.57E+01 | 2.77E-53 | BST2 | 3.13E+01 | 3.42E-213 | EIF2AK2 | 2.54E+01 | 7.75E-141 |
| 18 | TNFSF10 | 3.11E+01 | 2.21E-210 | STAT1 | 1.79E+01 | 2.17E-70 | ETV7 | 3.18E+01 | 2.04E-219 | EIF2AK2 | 1.47E+01 | 6.37E-47 | CASP1 | 3.13E+01 | 1.46E-212 | CASP1 | 2.46E+01 | 1.22E-131 |
| 19 | CASP1 | 2.95E+01 | 1.27E-189 | PARP9 | 1.79E+01 | 2.27E-70 | EIF2AK2 | 3.13E+01 | 8.60E-213 | TGM2 | 1.46E+01 | 3.19E-46 | NMI | 3.09E+01 | 1.36E-207 | CASP7 | 2.40E+01 | 5.34E-125 |
| 20 | IFITM3 | 2.93E+01 | 7.73E-187 | FHL2 | 1.76E+01 | 4.06E-68 | OPTN | 3.08E+01 | 1.03E-206 | STAT1 | 1.45E+01 | 1.98E-45 | IFITM3 | 2.97E+01 | 2.90E-191 | STAT1 | 2.37E+01 | 4.23E-122 |
| 21 | CASP7 | 2.75E+01 | 1.05E-164 | TGM2 | 1.75E+01 | 3.48E-67 | STAT1 | 3.08E+01 | 1.30E-206 | BCL10 | 1.43E+01 | 1.29E-44 | OPTN | 2.82E+01 | 2.00E-173 | OPTN | 2.28E+01 | 1.58E-113 |
| 22 | BIRC3 | 2.63E+01 | 2.45E-150 | CASP1 | 1.69E+01 | 6.99E-63 | TNFSF10 | 2.78E+01 | 1.50E-168 | PARK7 | 1.35E+01 | 8.16E-40 | ETV7 | 2.81E+01 | 3.91E-172 | ETV7 | 2.26E+01 | 1.19E-111 |
| 23 | ETV7 | 2.60E+01 | 1.91E-147 | ETV7 | 1.66E+01 | 1.22E-60 | CASP1 | 2.50E+01 | 2.79E-136 | HIF1A | 1.35E+01 | 1.31E-39 | STAT1 | 2.72E+01 | 1.48E-161 | TRIM38 | 2.23E+01 | 4.98E-108 |
| 24 | NMI | 2.56E+01 | 1.15E-142 | TLE4 | 1.62E+01 | 1.72E-57 | TGM2 | 2.28E+01 | 1.11E-112 | CFB | 1.34E+01 | 5.41E-39 | TMEM9B | 2.61E+01 | 9.91E-148 | NMI | 2.22E+01 | 1.58E-107 |
| 25 | CCAR1 | 2.46E+01 | 1.04E-131 | IFITM3 | 1.61E+01 | 1.49E-56 | BIRC3 | 2.25E+01 | 7.60E-110 | C3 | 1.34E+01 | 7.78E-39 | CASP7 | 2.49E+01 | 2.32E-135 | TMEM9B | 2.19E+01 | 9.84E-105 |

**Top 25 inactivated proteins**

| | Ciliated (3dpi) | | | Ciliated (6dpi) | | | Secretory (3dpi) | | | Secretory (6dpi) | | | Basal (3dpi) | | | Basal (6dpi) | | |
|---|---|---|---|---|---|---|---|---|---|---|---|---|---|---|---|---|---|---|
| rank | protein | NES | FDR | protein | NES | FDR | protein | NES | FDR | protein | NES | FDR | protein | NES | FDR | protein | NES | FDR |
| 1 | CAPS | -2.26E+01 | 5.98E-111 | SYTL3 | -2.47E+01 | 3.76E-131 | RPS6 | -3.51E+01 | 2.44E-268 | RPS6 | -1.67E+01 | 2.07E-60 | RPS6 | -3.23E+01 | 5.17E-227 | UBA52 | -2.84E+01 | 1.75E-175 |
| 2 | SOX5 | -2.24E+01 | 4.91E-109 | SLC22A4 | -2.46E+01 | 4.70E-130 | NACA | -2.71E+01 | 6.65E-160 | TOB1 | -1.47E+01 | 5.00E-47 | RPS3 | -2.96E+01 | 2.20E-190 | RPS6 | -2.84E+01 | 2.92E-175 |
| 3 | ZNF491 | -2.24E+01 | 7.00E-109 | CDHR3 | -2.43E+01 | 1.33E-127 | RPS3 | -2.48E+01 | 5.87E-134 | DUSP2 | -1.42E+01 | 4.20E-44 | NACA | -2.66E+01 | 3.65E-154 | RPS3 | -2.55E+01 | 7.42E-142 |
| 4 | PROS1 | -2.19E+01 | 1.30E-104 | ZNF440 | -2.37E+01 | 7.36E-122 | RPL7 | -2.27E+01 | 4.63E-112 | NACA | -1.38E+01 | 1.78E-41 | UBA52 | -2.56E+01 | 2.70E-142 | NACA | -2.52E+01 | 9.88E-138 |
| 5 | ESRRG | -2.15E+01 | 1.68E-100 | IFT172 | -2.35E+01 | 9.38E-120 | YBX1 | -2.26E+01 | 4.62E-111 | HIPK1 | -1.32E+01 | 4.13E-38 | YBX1 | -2.33E+01 | 1.10E-118 | POU2AF1 | -2.08E+01 | 1.49E-94 |
| 6 | RGS22 | -2.09E+01 | 1.18E-95 | ZNF491 | -2.35E+01 | 1.84E-119 | ZNF599 | -1.94E+01 | 5.97E-82 | FOS | -1.32E+01 | 7.48E-38 | RPL7 | -2.32E+01 | 1.10E-117 | HES5 | -2.03E+01 | 7.10E-90 |
| 7 | CERKL | -2.08E+01 | 8.11E-95 | STX2 | -2.35E+01 | 1.86E-119 | NPHP1 | -1.87E+01 | 1.30E-76 | GADD45B | -1.31E+01 | 1.48E-37 | PRMT1 | -1.81E+01 | 2.75E-71 | RPL7 | -2.02E+01 | 5.23E-89 |
| 8 | SYTL3 | -2.06E+01 | 7.38E-93 | HIPK1 | -2.34E+01 | 8.15E-119 | TRIM32 | -1.84E+01 | 5.00E-74 | RPS3 | -1.28E+01 | 1.82E-35 | POU2AF1 | -1.72E+01 | 5.82E-65 | YBX1 | -1.93E+01 | 3.49E-81 |
| 9 | ZNF157 | -2.01E+01 | 1.26E-88 | MAK | -2.31E+01 | 8.12E-116 | UBA52 | -1.83E+01 | 2.63E-73 | JADE1 | -1.27E+01 | 5.45E-35 | MAK | -1.67E+01 | 4.27E-61 | SP5 | -1.90E+01 | 1.54E-78 |
| 10 | OSBPL6 | -2.01E+01 | 1.91E-88 | ZNF19 | -2.31E+01 | 8.12E-116 | STOML3 | -1.82E+01 | 6.46E-72 | SIX4 | -1.26E+01 | 1.73E-34 | PHB2 | -1.60E+01 | 1.56E-56 | CATIP | -1.89E+01 | 6.67E-78 |
| 11 | ZNF19 | -2.01E+01 | 2.23E-88 | TRIP13 | -2.30E+01 | 4.81E-115 | LZTFL1 | -1.70E+01 | 4.18E-63 | TRIM32 | -1.23E+01 | 2.70E-33 | STX2 | -1.60E+01 | 4.77E-56 | UNCX | -1.87E+01 | 2.52E-76 |
| 12 | IL5RA | -2.01E+01 | 3.84E-88 | NEK11 | -2.30E+01 | 4.81E-115 | RFX3 | -1.64E+01 | 1.81E-58 | SLC22A4 | -1.22E+01 | 1.85E-32 | DLX4 | -1.55E+01 | 5.01E-53 | ARHGAP39 | -1.86E+01 | 1.97E-75 |
| 13 | SLC22A4 | -2.00E+01 | 1.83E-87 | DTHD1 | -2.28E+01 | 1.91E-112 | LMCD1 | -1.64E+01 | 1.87E-58 | TP53BP1 | -1.21E+01 | 5.27E-32 | ALX4 | -1.54E+01 | 2.56E-52 | ZFHX2 | -1.84E+01 | 3.50E-74 |
| 14 | STX2 | -1.92E+01 | 6.27E-81 | IFT57 | -2.26E+01 | 3.19E-111 | THAP10 | -1.63E+01 | 5.69E-58 | BCL9 | -1.20E+01 | 9.91E-32 | ZBBX | -1.54E+01 | 3.61E-52 | STX2 | -1.79E+01 | 4.78E-70 |
| 15 | SHANK2 | -1.89E+01 | 2.43E-78 | TP53BP1 | -2.23E+01 | 1.94E-108 | FOS | -1.63E+01 | 5.69E-58 | MOK | -1.17E+01 | 9.40E-30 | STOML3 | -1.52E+01 | 4.53E-51 | UTF1 | -1.79E+01 | 1.21E-69 |
| 16 | MS4A8 | -1.89E+01 | 8.49E-78 | JAZF1 | -2.18E+01 | 9.13E-104 | TUSC3 | -1.58E+01 | 1.18E-54 | JUNB | -1.16E+01 | 1.03E-29 | ZNF440 | -1.44E+01 | 1.08E-45 | LMCD1 | -1.78E+01 | 1.46E-69 |
| 17 | JADE1 | -1.88E+01 | 1.25E-77 | ZNF157 | -2.15E+01 | 8.21E-101 | EEF2 | -1.58E+01 | 1.32E-54 | UBA52 | -1.16E+01 | 1.61E-29 | EEF2 | -1.41E+01 | 4.58E-44 | DLX4 | -1.78E+01 | 3.22E-69 |
| 18 | LMCD1 | -1.86E+01 | 7.16E-76 | ZNF3 | -2.12E+01 | 8.80E-98 | ZNF440 | -1.57E+01 | 5.01E-54 | RORC | -1.15E+01 | 3.25E-29 | ZFP90 | -1.41E+01 | 4.93E-44 | STMN1 | -1.76E+01 | 9.95E-68 |
| 19 | ARHGAP18 | -1.77E+01 | 7.13E-69 | MAP3K19 | -2.09E+01 | 6.05E-95 | SLC22A4 | -1.54E+01 | 3.19E-52 | CERKL | -1.15E+01 | 6.59E-29 | DTHD1 | -1.39E+01 | 1.66E-42 | SLC22A4 | -1.76E+01 | 2.15E-67 |
| 20 | GLIS3 | -1.76E+01 | 7.42E-68 | GLIS3 | -2.07E+01 | 2.83E-93 | ZNF487 | -1.53E+01 | 1.63E-51 | ZFPM1 | -1.12E+01 | 1.98E-27 | HES5 | -1.38E+01 | 2.58E-42 | ALX4 | -1.76E+01 | 2.15E-67 |
| 21 | LZTFL1 | -1.76E+01 | 7.85E-68 | RP1 | -2.06E+01 | 3.51E-92 | DCDC1 | -1.52E+01 | 1.59E-50 | DUSP1 | -1.11E+01 | 3.16E-27 | SP5 | -1.38E+01 | 3.37E-42 | GSC2 | -1.75E+01 | 6.16E-67 |
| 22 | POU2AF1 | -1.74E+01 | 1.51E-66 | ZMYND12 | -2.06E+01 | 5.62E-92 | TRIP13 | -1.52E+01 | 2.07E-50 | THRA | -1.11E+01 | 3.90E-27 | SLC22A4 | -1.38E+01 | 7.30E-42 | SIM2 | -1.74E+01 | 2.83E-66 |
| 23 | ZNF599 | -1.71E+01 | 4.21E-64 | NPHP1 | -2.02E+01 | 5.48E-89 | HIPK1 | -1.50E+01 | 2.06E-49 | SHANK2 | -1.11E+01 | 4.97E-27 | RP1 | -1.36E+01 | 4.00E-41 | CDHR3 | -1.74E+01 | 3.27E-66 |
| 24 | IFT57 | -1.65E+01 | 1.61E-59 | DCDC1 | -2.00E+01 | 5.24E-87 | TP53BP1 | -1.50E+01 | 2.51E-49 | RPL7 | -1.10E+01 | 7.36E-27 | RUVBL1 | -1.36E+01 | 5.83E-41 | DMRTC2 | -1.73E+01 | 1.05E-65 |
| 25 | JAZF1 | -16.405241 | 3.72E-59 | SIX4 | -19.904005 | 2.22E-86 | ZBTB4 | -14.802814 | 4.32E-48 | NEK11 | -10.874796 | 4.68E-26 | TCEA2 | -13.498491 | 2.32E-40 | RAB36 | -17.311389 | 1.20E-65 |

### Supplementary Table 3b

Top Activated and inactivated MRs in the host response signatures **Infected vs Bystander**

| Top 25 activated proteins | | | | | | | | | |
|---|---|---|---|---|---|---|---|---|---|
| | Ciliated (3dpi) | | | Secretory (3dpi) | | | Basal (3dpi) | | |
| rank | protein | NES | FDR | protein | NES | FDR | protein | NES | FDR |
| 1 | ITGA2 | 2.07E+01 | 7.93E-92 | KAT2B | 1.31E+01 | 1.24E-35 | ZNF143 | 1.21E+01 | 3.75E-30 |
| 2 | NCOR1 | 1.79E+01 | 2.63E-68 | SMC3 | 1.28E+01 | 2.40E-34 | UBA3 | 1.20E+01 | 4.58E-30 |
| 3 | HDAC1 | 1.76E+01 | 3.12E-66 | LIN7C | 1.11E+01 | 5.42E-26 | MED7 | 1.16E+01 | 2.81E-28 |
| 4 | EHBP1 | 1.71E+01 | 1.32E-62 | APEX1 | 1.01E+01 | 2.33E-21 | MDH2 | 1.15E+01 | 8.98E-28 |
| 5 | MTCH1 | 1.51E+01 | 3.43E-49 | TRIP11 | 9.85E+00 | 2.12E-20 | RAN | 1.11E+01 | 6.89E-26 |
| 6 | ZNF567 | 1.50E+01 | 2.02E-48 | SMAD4 | 9.59E+00 | 2.52E-19 | HSBP1 | 1.04E+01 | 9.70E-23 |
| 7 | RBFOX2 | 1.47E+01 | 2.20E-46 | ZNF451 | 9.50E+00 | 5.07E-19 | GABPB1 | 1.02E+01 | 8.16E-22 |
| 8 | SMARCC1 | 1.47E+01 | 2.47E-46 | PRKAR2A | 9.36E+00 | 1.73E-18 | SHOC2 | 1.01E+01 | 9.69E-22 |
| 9 | STIM2 | 1.44E+01 | 1.60E-44 | MCMBP | 9.34E+00 | 1.99E-18 | PPP2R1A | 9.65E+00 | 1.01E-19 |
| 10 | MED21 | 1.43E+01 | 8.97E-44 | XRCC5 | 8.58E+00 | 1.49E-15 | CCNC | 9.60E+00 | 1.49E-19 |
| 11 | ZNF280C | 1.37E+01 | 1.54E-40 | WWTR1 | 8.56E+00 | 1.58E-15 | SAP18 | 9.44E+00 | 5.83E-19 |
| 12 | ADNP2 | 1.37E+01 | 2.12E-40 | RAD1 | 8.45E+00 | 3.67E-15 | CAPNS1 | 9.16E+00 | 7.05E-18 |
| 13 | PPP2R5A | 1.37E+01 | 2.42E-40 | HDAC1 | 8.44E+00 | 3.79E-15 | UBQLN2 | 9.07E+00 | 1.59E-17 |
| 14 | ANK3 | 1.36E+01 | 5.68E-40 | DSG2 | 8.33E+00 | 8.21E-15 | KDM4B | 8.98E+00 | 3.16E-17 |
| 15 | CREB1 | 1.35E+01 | 1.53E-39 | STX7 | 8.14E+00 | 3.73E-14 | FOXN2 | 8.96E+00 | 3.88E-17 |
| 16 | KMT2C | 1.32E+01 | 6.62E-38 | RAB3GAP2 | 8.13E+00 | 3.86E-14 | SNAP23 | 8.88E+00 | 7.41E-17 |
| 17 | HMBOX1 | 1.32E+01 | 9.08E-38 | NOTCH3 | 8.04E+00 | 7.64E-14 | DPF2 | 8.86E+00 | 7.85E-17 |
| 18 | USP33 | 1.31E+01 | 2.52E-37 | IKZF5 | 8.01E+00 | 9.57E-14 | SMARCA5 | 8.86E+00 | 7.85E-17 |
| 19 | BLZF1 | 1.31E+01 | 4.95E-37 | SRP72 | 8.00E+00 | 9.85E-14 | RAB3GAP2 | 8.77E+00 | 1.75E-16 |
| 20 | BPTF | 1.31E+01 | 4.95E-37 | PTPN2 | 7.91E+00 | 1.90E-13 | DYNC1LI1 | 8.61E+00 | 6.75E-16 |
| 21 | ZBTB38 | 1.30E+01 | 1.35E-36 | UBA3 | 7.88E+00 | 2.38E-13 | NAA15 | 8.54E+00 | 1.14E-15 |
| 22 | SNAPC3 | 1.29E+01 | 2.73E-36 | ZCCHC17 | 7.86E+00 | 2.55E-13 | ILF2 | 8.45E+00 | 2.49E-15 |
| 23 | CEBPZ | 1.26E+01 | 1.17E-34 | PCBD1 | 7.78E+00 | 4.95E-13 | USP14 | 8.25E+00 | 1.28E-14 |
| 24 | MACC1 | 1.26E+01 | 2.12E-34 | THAP2 | 7.67E+00 | 1.09E-12 | TANK | 8.19E+00 | 2.02E-14 |
| 25 | PPP1R13L | 1.26E+01 | 2.70E-34 | CBX4 | 7.60E+00 | 1.71E-12 | HDAC2 | 8.14E+00 | 2.94E-14 |

| Top 25 inactivated proteins | | | | | | | | | |
|---|---|---|---|---|---|---|---|---|---|
| | Ciliated (3dpi) | | | Secretory (3dpi) | | | Basal (3dpi) | | |
| rank | protein | NES | FDR | protein | NES | FDR | protein | NES | FDR |
| 1 | NACC2 | -1.17E+01 | 6.68E-30 | ELF3 | -1.13E+01 | 6.67E-27 | NUCKS1 | -1.20E+01 | 4.58E-30 |
| 2 | CAPS | -1.14E+01 | 1.59E-28 | HDAC7 | -1.11E+01 | 8.69E-26 | ZNF746 | -9.66E+00 | 9.47E-20 |
| 3 | LYL1 | -1.13E+01 | 5.07E-28 | NCOA3 | -9.24E+00 | 4.54E-18 | KIFAP3 | -9.46E+00 | 5.32E-19 |
| 4 | MARK2 | -1.12E+01 | 1.74E-27 | ERBB2 | -8.95E+00 | 6.20E-17 | EBF4 | -9.25E+00 | 3.28E-18 |
| 5 | PTK2 | -1.12E+01 | 2.98E-27 | SCML1 | -8.53E+00 | 2.01E-15 | MED31 | -8.25E+00 | 1.28E-14 |
| 6 | MEF2D | -1.10E+01 | 2.00E-26 | NPAS3 | -8.44E+00 | 3.71E-15 | ZNF740 | -7.37E+00 | 6.90E-12 |
| 7 | IFT52 | -1.09E+01 | 3.03E-26 | ARL6 | -8.41E+00 | 4.41E-15 | HES4 | -7.33E+00 | 9.23E-12 |
| 8 | DNER | -1.08E+01 | 1.31E-25 | CARF | -8.28E+00 | 1.22E-14 | ZBTB46 | -7.03E+00 | 7.09E-11 |
| 9 | ZFP41 | -1.08E+01 | 1.69E-25 | CAPN1 | -8.24E+00 | 1.58E-14 | TCEA2 | -6.93E+00 | 1.36E-10 |
| 10 | DUSP2 | -1.07E+01 | 2.05E-25 | ULK1 | -7.90E+00 | 2.09E-13 | ZNF438 | -6.82E+00 | 2.71E-10 |
| 11 | ARF3 | -1.04E+01 | 5.37E-24 | PCBP1 | -7.70E+00 | 8.73E-13 | LZTFL1 | -6.78E+00 | 3.53E-10 |
| 12 | PCBP1 | -1.04E+01 | 6.84E-24 | MXD4 | -7.66E+00 | 1.09E-12 | WDPCP | -6.77E+00 | 3.69E-10 |
| 13 | STAT3 | -1.02E+01 | 4.06E-23 | NECTIN4 | -7.66E+00 | 1.12E-12 | ZFP41 | -6.76E+00 | 3.93E-10 |
| 14 | ZBTB48 | -1.02E+01 | 4.59E-23 | ITGB6 | -7.55E+00 | 2.41E-12 | CD81 | -6.75E+00 | 3.97E-10 |
| 15 | ZDHHC5 | -1.01E+01 | 1.27E-22 | MTA2 | -7.48E+00 | 3.44E-12 | CDC42BPB | -6.70E+00 | 5.59E-10 |
| 16 | CCT3 | -1.01E+01 | 1.46E-22 | USP22 | -7.48E+00 | 3.44E-12 | SRI | -6.68E+00 | 6.16E-10 |
| 17 | TGM2 | -1.01E+01 | 1.59E-22 | EHMT1 | -7.38E+00 | 6.73E-12 | POU6F1 | -6.67E+00 | 6.63E-10 |
| 18 | EHF | -9.87E+00 | 1.27E-21 | ERBB3 | -7.23E+00 | 1.84E-11 | BARX1 | -6.66E+00 | 6.97E-10 |
| 19 | ASXL2 | -9.83E+00 | 1.86E-21 | ZBTB46 | -7.22E+00 | 1.87E-11 | IRAK1BP1 | -6.66E+00 | 6.97E-10 |
| 20 | NRIP1 | -9.79E+00 | 2.71E-21 | ZNF287 | -7.12E+00 | 3.91E-11 | MUC15 | -6.64E+00 | 7.84E-10 |
| 21 | CIAO1 | -9.60E+00 | 1.53E-20 | FCHO2 | -6.95E+00 | 1.21E-10 | PCYOX1 | -6.61E+00 | 9.52E-10 |
| 22 | PARP1 | -9.58E+00 | 1.93E-20 | RAB25 | -6.89E+00 | 1.83E-10 | ZNF487 | -6.52E+00 | 1.68E-09 |
| 23 | FOXP4 | -9.44E+00 | 6.51E-20 | ITPR3 | -6.80E+00 | 3.19E-10 | MARK3 | -6.40E+00 | 3.51E-09 |
| 24 | JPT2 | -9.23E+00 | 4.33E-19 | STAT5A | -6.80E+00 | 3.27E-10 | ZNF226 | -6.35E+00 | 4.61E-09 |
| 25 | IFI27 | -9.22E+00 | 4.96E-19 | CNOT8 | -6.78E+00 | 3.54E-10 | CHD3 | -6.28E+00 | 7.07E-09 |

### Supplementary Table 4

#### Top Activated and inactivated MR of drugs predicted by ViroTreat

##### Top 25 Activated MRs of 11 drugs predicted by ViroTreat

| Drug | Bedaquiline | | Lomustine | | Ethinyl Estradiol | | Tolperisone HCl | | Vonoprazan Fumarate | | Oclacitinib maleate | |
|---|---|---|---|---|---|---|---|---|---|---|---|---|
| Rank | protein | NES | protein | NES | protein | NES | protein | NES | protein | NES | protein | NES |
| 1 | CBX4 | 2.50 | PRRX2 | 2.79 | EREG | 3.22 | AIRE | 2.96 | HYAL2 | 4.34 | ZNF653 | 2.90 |
| 2 | IRF3 | 2.48 | PQBP1 | 2.58 | GPRC5A | 3.19 | HNF4A | 2.83 | AIRE | 4.02 | MAMSTR | 2.66 |
| 3 | RPS3 | 2.45 | LTBR | 2.39 | FHL2 | 2.97 | DMRTC2 | 2.82 | LMX1A | 3.91 | HES5 | 2.45 |
| 4 | EEF2 | 2.45 | CUX1 | 2.33 | GRHL3 | 2.92 | ONECUT1 | 2.76 | LYL1 | 3.79 | TGFB1I1 | 2.33 |
| 5 | ATP5F1A | 2.40 | IRF7 | 2.25 | PRRX2 | 2.73 | DMRTB1 | 2.73 | CBFA2T3 | 3.79 | UNCX | 2.32 |
| 6 | ZNF446 | 2.37 | ELK1 | 2.17 | ZNF446 | 2.66 | POU2F2 | 2.72 | VSX2 | 3.74 | OLIG1 | 2.28 |
| 7 | CCNE1 | 2.29 | ZNF219 | 2.16 | IGFBP6 | 2.63 | CBFA2T3 | 2.71 | POU3F2 | 3.73 | FEZF1 | 2.24 |
| 8 | ZMYND10 | 2.27 | PBXIP1 | 2.16 | FEZF1 | 2.46 | KLF13 | 2.68 | HNF4A | 3.71 | CBFA2T3 | 2.17 |
| 9 | CTBP2 | 2.25 | NFATC2IP | 2.12 | AIRE | 2.44 | VAX2 | 2.67 | CCNE1 | 3.71 | POU3F3 | 2.16 |
| 10 | CITED4 | 2.17 | CRB3 | 2.08 | HR | 2.43 | TBX10 | 2.64 | SIM2 | 3.69 | ELANE | 2.13 |
| 11 | E4F1 | 2.14 | RAMP1 | 2.03 | CRB3 | 2.33 | FOXG1 | 2.63 | NFATC4 | 3.69 | GATA5 | 2.12 |
| 12 | PLEC | 1.97 | HMG20B | 2.02 | HYAL2 | 2.31 | SLC39A4 | 2.61 | GSC2 | 3.67 | TLE6 | 2.09 |
| 13 | PCGF2 | 1.91 | FEZF1 | 2.02 | UNCX | 2.27 | ZNF488 | 2.58 | BHLHE23 | 3.63 | MAP3K10 | 2.07 |
| 14 | HMX3 | 1.89 | CDKN1C | 1.99 | CFB | 2.27 | KLF1 | 2.57 | HES5 | 3.63 | SLC22A4 | 2.06 |
| 15 | RPS6 | 1.87 | ZFP41 | 1.98 | FGFBP1 | 2.26 | PITX2 | 2.56 | DMRTC2 | 3.59 | MYPOP | 2.02 |
| 16 | UBA52 | 1.87 | THRA | 1.96 | BAX | 2.24 | TCF23 | 2.52 | TFAP2E | 3.56 | TBX2 | 2.00 |
| 17 | ROPN1L | 1.86 | BAD | 1.96 | TONSL | 2.22 | LMX1A | 2.50 | TBX10 | 3.53 | NACC2 | 1.99 |
| 18 | AIRE | 1.76 | GIPC1 | 1.95 | PAX4 | 2.22 | GSX2 | 2.48 | FGF2 | 3.53 | TONSL | 1.96 |
| 19 | MAZ | 1.74 | TCF15 | 1.94 | HNF4A | 2.20 | HOXD9 | 2.47 | TLX2 | 3.50 | SP4 | 1.96 |
| 20 | PRDM16 | 1.74 | SLC2A4RG | 1.94 | LMX1A | 2.16 | POU3F2 | 2.46 | MAMSTR | 3.48 | ONECUT1 | 1.92 |
| 21 | HNF4A | 1.74 | FOXE3 | 1.93 | ZBTB7B | 2.15 | VSX2 | 2.45 | STMN1 | 3.47 | EGR4 | 1.91 |
| 22 | PRMT1 | 1.72 | PLEC | 1.88 | BSX | 2.15 | FGF2 | 2.35 | ACTL6B | 3.42 | AIRE | 1.90 |
| 23 | LMCD1 | 1.70 | SOX12 | 1.87 | MAFB | 2.14 | STMN1 | 2.34 | ISX | 3.41 | FHL3 | 1.89 |
| 24 | PRDM13 | 1.69 | DEF8 | 1.87 | TBX10 | 2.14 | PRDM13 | 2.32 | ZFPM1 | 3.40 | ONECUT2 | 1.88 |
| 25 | TBX2 | 1.68 | MNX1 | 1.87 | ZNF488 | 2.14 | EGR4 | 2.32 | DDN | 3.40 | PRDM16 | 1.87 |

| Drug | Stearic acid | | Anisodamine Hydrobromide | | Prasugrel | | Aprepitant | | Dutasteride | |
|---|---|---|---|---|---|---|---|---|---|---|
| Rank | protein | NES | protein | NES | protein | NES | protein | NES | protein | NES |
| 1 | TRIB3 | 3.22 | MAMSTR | 2.48 | TRIB3 | 3.06 | EID2 | 2.53 | SMYD1 | 2.50 |
| 2 | HES5 | 3.17 | PROP1 | 2.23 | SLC39A4 | 2.64 | ADM | 2.52 | CDKN1C | 2.25 |
| 3 | SP5 | 3.09 | VSX2 | 2.23 | EGR4 | 2.60 | TCF15 | 2.46 | DUSP2 | 2.25 |
| 4 | TONSL | 3.00 | ZNF488 | 2.15 | DUSP2 | 2.57 | SOX12 | 2.45 | DMRTC2 | 2.13 |
| 5 | TGFB1I1 | 2.99 | GATA5 | 2.12 | EREG | 2.42 | ZNF444 | 2.44 | ZNF366 | 2.13 |
| 6 | ZFPM1 | 2.96 | LMX1A | 2.07 | MAFF | 2.41 | GSX2 | 2.33 | GSX2 | 2.06 |
| 7 | PROP1 | 2.93 | CDKN1C | 2.07 | ZNF444 | 2.40 | UBA52 | 2.32 | PRRX2 | 2.05 |
| 8 | POU3F3 | 2.86 | HOXD13 | 2.02 | KLF1 | 2.39 | DMRTB1 | 2.30 | KCNK1 | 2.02 |
| 9 | FOXI2 | 2.85 | CBFA2T3 | 2.00 | ZNF628 | 2.38 | BSX | 2.30 | TLX2 | 1.99 |
| 10 | UNCX | 2.81 | DDN | 1.99 | HIC1 | 2.35 | KLF1 | 2.28 | PRDM12 | 1.97 |
| 11 | HMX3 | 2.77 | DTX1 | 1.96 | GRHL3 | 2.34 | EBF1 | 2.25 | TRIB3 | 1.96 |
| 12 | DLX4 | 2.75 | HNF1A | 1.94 | PAX2 | 2.34 | SLC39A4 | 2.20 | CITED4 | 1.94 |
| 13 | TLX2 | 2.74 | OLIG1 | 1.88 | DMRTC2 | 2.33 | TCF24 | 2.18 | KLF2 | 1.93 |
| 14 | GSC2 | 2.74 | SMYD1 | 1.88 | VSX2 | 2.29 | SMYD1 | 2.13 | ZBTB7B | 1.93 |
| 15 | BSX | 2.72 | DRGX | 1.88 | PIM1 | 2.22 | ZNF488 | 2.13 | MNX1 | 1.92 |
| 16 | NR2E1 | 2.72 | ANKRD1 | 1.88 | NR2E1 | 2.22 | NFE2 | 2.11 | FOXE3 | 1.91 |
| 17 | GATA5 | 2.70 | STMN1 | 1.87 | CD151 | 2.20 | MAZ | 2.10 | ETV3L | 1.88 |
| 18 | MAMSTR | 2.68 | NAMPT | 1.86 | VAX2 | 2.18 | ELANE | 2.10 | NACC2 | 1.87 |
| 19 | GFI1 | 2.68 | LHX1 | 1.84 | TCEAL3 | 2.17 | NEUROD2 | 2.10 | ISX | 1.84 |
| 20 | BARX1 | 2.67 | PRDM16 | 1.83 | ACTL6B | 2.16 | SIX5 | 2.08 | IGFBP6 | 1.82 |
| 21 | PRDM16 | 2.64 | POU2F2 | 1.83 | SOCS3 | 2.13 | ZBTB7B | 2.08 | LMX1A | 1.75 |
| 22 | EGR4 | 2.62 | KLF1 | 1.81 | BSX | 2.13 | SLC6A8 | 2.06 | NFE2 | 1.72 |
| 23 | PURG | 2.60 | ZFPM1 | 1.80 | SOX18 | 2.09 | AIRE | 2.06 | DLX2 | 1.69 |
| 24 | MED9 | 2.59 | HNF4A | 1.77 | HYAL2 | 2.08 | FEZF1 | 2.05 | VAX2 | 1.69 |
| 25 | MYPOP | 2.58 | EGR4 | 1.77 | SOX11 | 2.08 | CBFA2T3 | 2.05 | NEUROD4 | 1.69 |

##### Top 25 Inactivated MRs of 11 drugs predicted by ViroTreat

| Drug | Bedaquiline | | Lomustine | | Ethinyl Estradiol | | Tolperisone HCl | | Vonoprazan Fumarate | | Oclacitinib maleate | |
|---|---|---|---|---|---|---|---|---|---|---|---|---|
| Rank | protein | NES | protein | NES | protein | NES | protein | NES | protein | NES | protein | NES |
| 1 | FUBP1 | -2.82 | RYBP | -3.08 | N4BP2L2 | -3.54 | ZNF487 | -3.01 | RLIM | -4.32 | ZMYM5 | -2.58 |
| 2 | PRKAA1 | -2.81 | ITCH | -2.58 | DMTF1 | -3.15 | ARL6 | -2.77 | IFT88 | -4.10 | ZNF92 | -2.51 |
| 3 | MED4 | -2.80 | DOCK1 | -2.54 | ZRANB2 | -3.09 | TBPL1 | -2.75 | LZTFL1 | -4.09 | SETD3 | -2.41 |
| 4 | MTA3 | -2.73 | DLG1 | -2.51 | TUT4 | -3.08 | ZNF273 | -2.71 | PSMA4 | -3.98 | PIK3C2A | -2.39 |
| 5 | ZNF322 | -2.71 | PFDN1 | -2.48 | SUPT20H | -3.00 | WDPCP | -2.56 | CDK7 | -3.94 | RAB14 | -2.39 |
| 6 | NFXL1 | -2.70 | OGT | -2.42 | ZMYM4 | -2.99 | CEP290 | -2.56 | GDI2 | -3.83 | RLIM | -2.18 |
| 7 | ATRX | -2.67 | ELF1 | -2.36 | GPR162 | -2.92 | EVI5 | -2.54 | BIRC2 | -3.82 | GSTP1 | -2.18 |
| 8 | STXBP3 | -2.66 | SMAD1 | -2.34 | ZNF292 | -2.90 | KIFAP3 | -2.50 | MED7 | -3.82 | IKZF5 | -2.16 |
| 9 | USP33 | -2.66 | CHD6 | -2.32 | DNAAF1 | -2.84 | ZCCHC17 | -2.48 | NRIP1 | -3.80 | ZNF639 | -2.15 |
| 10 | CUL5 | -2.65 | FOXN2 | -2.32 | CARF | -2.83 | ATRX | -2.47 | ZFP3 | -3.69 | ADNP2 | -2.09 |
| 11 | CHD1 | -2.44 | N4BP2L2 | -2.27 | CDHR3 | -2.78 | ZNF391 | -2.44 | IFT57 | -3.68 | SMAD2 | -2.06 |
| 12 | ZNF189 | -2.44 | DMTF1 | -2.19 | SUPT7L | -2.77 | DCDC1 | -2.43 | COPS5 | -3.67 | GTF2A1 | -2.06 |
| 13 | MAP4K5 | -2.39 | CREB1 | -2.18 | RFX2 | -2.77 | SMC3 | -2.42 | SPA17 | -3.64 | ZNF24 | -2.04 |
| 14 | CSNK1G3 | -2.38 | BRWD1 | -2.18 | OGT | -2.72 | ROCK1 | -2.42 | ARL6 | -3.59 | PEX11B | -2.03 |
| 15 | CD47 | -2.36 | ZMYM4 | -2.16 | WWC1 | -2.72 | ZFC3H1 | -2.39 | SOD1 | -3.58 | SRP72 | -2.02 |
| 16 | LCOR | -2.30 | COG3 | -2.16 | PHIP | -2.68 | TRIP11 | -2.36 | THRAP3 | -3.54 | UTP11 | -2.00 |
| 17 | NFYB | -2.28 | ZNF585A | -2.16 | NSD3 | -2.63 | ZNF599 | -2.34 | GTF2H5 | -3.53 | SUMO1 | -1.98 |
| 18 | MED7 | -2.27 | USP14 | -2.13 | RAB36 | -2.61 | TIAL1 | -2.33 | ZBBX | -3.50 | SMAD5 | -1.97 |
| 19 | CEBPZ | -2.26 | MED14 | -2.11 | ZNF644 | -2.59 | CNOT4 | -2.32 | GTF2A2 | -3.50 | BCLAF1 | -1.97 |
| 20 | PCGF3 | -2.15 | ZNF292 | -2.11 | BASP1 | -2.58 | KDM3A | -2.30 | UBA52 | -3.49 | SLC31A1 | -1.97 |
| 21 | AFF1 | -2.13 | DNAJA1 | -2.11 | LCOR | -2.58 | ZBBX | -2.30 | SMARCA5 | -3.48 | STRAP | -1.97 |
| 22 | ZNF451 | -2.11 | WDR43 | -2.09 | PIAS1 | -2.57 | ATR | -2.28 | RAB28 | -3.46 | PCYOX1 | -1.96 |
| 23 | CASP8AP2 | -2.09 | RABL3 | -2.08 | SCAI | -2.56 | RFX3 | -2.26 | ZNF267 | -3.42 | CREBZF | -1.95 |
| 24 | CGGBP1 | -2.08 | GABPB1 | -2.07 | ZFHX2 | -2.55 | STX7 | -2.25 | IRAK1BP1 | -3.41 | KCNK1 | -1.94 |
| 25 | RFC1 | -2.07 | SUPT3H | -2.05 | MLLT10 | -2.52 | HNRNPK | -2.24 | YWHAE | -3.41 | COG3 | -1.93 |

| Drug | Stearic acid | | Anisodamine Hydrobromide | | Prasugrel | | Aprepitant | | Dutasteride | |
|---|---|---|---|---|---|---|---|---|---|---|
| Rank | protein | NES | protein | NES | protein | NES | protein | NES | protein | NES |
| 1 | EVI5 | -3.26 | CAPNS1 | -3.04 | ZNF287 | -2.60 | MTDH | -3.29 | GTF2H5 | -3.24 |
| 2 | NAE1 | -3.20 | RAB5C | -3.00 | ZFP3 | -2.59 | DNAJA1 | -3.02 | HSBP1 | -3.05 |
| 3 | ZMYM2 | -3.10 | UBQLN2 | -2.90 | ZFC3H1 | -2.47 | CIR1 | -2.74 | RAB36 | -2.97 |
| 4 | ELF2 | -3.09 | ADRM1 | -2.67 | SNX4 | -2.45 | SKIL | -2.73 | PSMC3IP | -2.54 |
| 5 | TDRD3 | -3.04 | TMEM9B | -2.38 | NR2C1 | -2.43 | PSMA4 | -2.68 | GPR162 | -2.53 |
| 6 | CEP89 | -3.03 | TMEM30B | -2.35 | WDPCP | -2.37 | RYBP | -2.67 | N4BP2L2 | -2.40 |
| 7 | ADNP2 | -2.98 | FUBP1 | -2.33 | ADNP | -2.35 | SERINC3 | -2.56 | RFX2 | -2.39 |
| 8 | METAP2 | -2.87 | MSRB2 | -2.32 | SMC3 | -2.32 | BASP1 | -2.56 | APEX1 | -2.33 |
| 9 | GTF2A1 | -2.86 | ARF5 | -2.31 | PATJ | -2.30 | ERBIN | -2.48 | DMTF1 | -2.26 |
| 10 | SS18 | -2.85 | NFE2L2 | -2.30 | ZMYM4 | -2.24 | CREB1 | -2.48 | OGT | -2.23 |
| 11 | CSNK1A1 | -2.84 | RABL3 | -2.29 | KLF12 | -2.23 | PRKDC | -2.45 | RPS6 | -2.23 |
| 12 | LZTFL1 | -2.83 | NCSTN | -2.29 | PRKDC | -2.23 | IQGAP1 | -2.45 | YWHAQ | -2.22 |
| 13 | SETD3 | -2.82 | PAWR | -2.29 | AEBP2 | -2.22 | OGT | -2.43 | GOPC | -2.21 |
| 14 | URI1 | -2.76 | DAG1 | -2.26 | PIAS1 | -2.22 | EVI5 | -2.35 | POLR2K | -2.18 |
| 15 | BTAF1 | -2.76 | HDAC3 | -2.21 | MED31 | -2.20 | ZSCAN16 | -2.35 | STYXL1 | -2.17 |
| 16 | COPS5 | -2.72 | FOXJ3 | -2.16 | ZC3H6 | -2.18 | ATP6AP2 | -2.34 | PARK7 | -2.16 |
| 17 | CNOT2 | -2.71 | AP2M1 | -2.11 | G3BP2 | -2.17 | IMPA1 | -2.34 | ZNF491 | -2.15 |
| 18 | BRD7 | -2.68 | FOXA1 | -2.06 | ZNF273 | -2.16 | UBE2L3 | -2.34 | CNOT4 | -2.13 |
| 19 | IKZF5 | -2.66 | RYBP | -2.04 | CREBL2 | -2.14 | PAFAH1B1 | -2.33 | SERINC3 | -2.11 |
| 20 | STAM | -2.64 | NME2 | -2.02 | ZNF138 | -2.13 | KDM1A | -2.32 | SPA17 | -2.10 |
| 21 | PIAS1 | -2.64 | ROCK1 | -2.01 | TBPL1 | -2.11 | ZNF322 | -2.32 | ZSCAN16 | -2.08 |
| 22 | COPS3 | -2.60 | MIER1 | -2.00 | HBP1 | -2.11 | PARP1 | -2.31 | PFDN1 | -2.06 |
| 23 | PIAS2 | -2.60 | TLE4 | -2.00 | TADA1 | -2.06 | N4BP2L2 | -2.31 | CDK7 | -2.05 |
| 24 | RICTOR | -2.60 | ADIPOR1 | -1.99 | ZSCAN16 | -1.98 | TAF11 | -2.29 | RYBP | -2.05 |
| 25 | RAB18 | -2.59 | ZNF189 | -1.99 | ZMYM5 | -1.97 | BRWD1 | -2.28 | CDK12 | -2.05 |
